# Skeletal muscle proteomic responses to energy deficit with concomitant aerobic exercise in humans

**DOI:** 10.1101/2025.05.15.654297

**Authors:** Y Nishimura, C Langan-Evans, HL Taylor, WL Foo, JP Morton, S Shepherd, J Strauss, JG Burniston, JL Areta

## Abstract

Energy deficit is a potent physiological stressor that has shaped human evolution and can improve lifespan and healthspan in a wide range of species. Preserving locomotive capacity was likely essential for survival during the human hunter-gatherer period but surprisingly little is known about the molecular effects of energy deficit on human skeletal muscle, which is a key tissue for locomotion and metabolic health. Here we show that a 5-day 78% reduction in energy availability with concomitant aerobic exercise in healthy men leads to a profound modulation of skeletal muscle phenotype alongside increases in fat oxidation at rest and during exercise and a 2.1 ±0.8 kg loss of fat free mass and 0.8 ±0.6 kg of fat mass. We used stable isotope (D_2_O) labelling and peptide mass spectrometry to investigate the abundance and turnover rates of individual proteins. Abundance (1469 proteins) and synthesis rate (736 proteins) data discovered a shift toward a more oxidative phenotype and reorganisation of cytoskeleton and extracellular matrix structure during energy deficit. Mitochondrial components: TCA, electron transport chain and beta-oxidation, were prominently represented amongst proteins that increased in abundance and synthesis rate, as well as proteins related to mitochondrial proteostasis, remodelling and quality-control such as BDH1 and LONP1. Changes in muscle metabolic pathways occurred alongside a reduction in extracellular matrix proteins, which may counteract the age-related muscle fibrosis. Our results suggest that muscle metabolic pathways are not only preserved but positively affected during periods of concomitant low energy availability and exercise.

## INTRODUCTION

During evolution humans may have adapted to endure periods of intermittent food availability alongside sustained needs for high levels of physical activity to procure food, shelter or evade danger (Prentice, 2005; J. Speakman, 2007). In the face of limited energy availability, life history models of energy allocation predict that energetic resources will be re-distributed among competing physiological processes in an order of priority that maximises survival (Pontzer & McGrosky, 2022). In the ancestorial human habitat (that of hunter-gatherers), suppression of growth and reproductive function would spare limited energetic resources during periods of food scarcity (Jasienska et al., 2017), whereas a reduction in physical capacity (necessary for food procurement) would have been less favourable (Jasienska, 2003). Given also the remarkable capacity for endurance in the genus Homo, possibly in relation to persistence hunting and scavenging (Bramble & Lieberman, 2004; Morin & Winterhalder, 2024), human physiology may have adapted to prioritise endurance capacity. Thus, the physiological machinery that supports locomotion may be preserved even during low energy availability, at the expense of other energetically demanding processes such as growth and reproductive function.

Energy prioritisation has far-reaching implications for several modern populations where energy deficit is either intentional or unavoidable. In healthy ageing, energy restriction is explored for its potential to extend lifespan and healthspan (Das et al., 2023; J. R. Speakman & Mitchell, 2011); in athletic populations, low energy availability is linked to adverse effects on health and performance (Areta et al., 2021; Jeukendrup et al., 2024; Van Rosmalen et al., 2024); in overweight or obese individuals, energy deficit underpins interventions aimed at improving metabolic health through weight loss (Magkos et al., 2016); and in space exploration, astronauts are reported to be in energy deficit (Bergouignan, 2016). A common and critical tissue in all these scenarios is skeletal muscle—essential for locomotion and central to metabolic regulation.

The crucial role of skeletal muscle has recently come to the forefront of public attention due to the growing use of GLP-1 receptor agonists, which are effective for weight loss, but can cause substantial muscle loss (Prado et al., 2024). It is not clear, however, whether muscle loss associated with GLP-1 receptor agonists is clinically relevant (Conte et al., 2024). A comprehensive analysis of how energy deficit affects skeletal muscle phenotype alongside the endocrine and physiological responses of energy preservation is yet to be conducted. An enhanced mechanistic understanding of the skeletal muscle phenotype shift with energy restriction and concomitant exercise in humans will not only identify key pathways modulated by energy deficit but also help to understand and predict the potential risks and benefits of interventions aiming to enhance muscle quality and function in the face of energy deficit.

Earlier studies that have investigated the effect of energy deficit on skeletal muscle in humans are limited to skeletal muscle bulk protein synthesis/breakdown (Areta et al., 2014; Caldwell et al., 2024; Carbone et al., 2014; Oxfeldt et al., 2023; Pasiakos et al., 2010), provide limited insights through addressing selected molecular markers (Coen et al., 2015; Rosenbaum et al., 2018; Toledo et al., 2006), or provided only insights at the gene expression level (Beals et al., 2023; Das et al., 2023; Yang et al., 2016). Protein turnover is a highly responsive characteristic of the muscle proteome that is required to maintain protein homeostasis and underpins the process of muscle adaptation. The processes of protein turnover (e.g. ribosomal translation, protein chaperoning and degradation via the ubiquitin proteasome system) are energetically costly and highly responsive to changes in muscle activity and feeding. Muscle proteins exhibit a wide range of different turnover rates and no study to date has investigated skeletal muscle proteome dynamics comprehensively in response to energy deficit. Dynamic proteomic profiling is a new technique built on the methodological developments in the fields of stable isotopic labelling, proteomics and computational biology, allowing to determine the synthesis, abundance and degradation human skeletal muscle proteins on a protein-by-protein basis (Burniston, 2019; Camera et al., 2017). Dynamic proteome profiling can characterise hundreds of skeletal muscle proteins to systematically and agnostically study tissue-specific protein dynamics (Hesketh et al., 2020; Nishimura et al., 2023), which may help determine the effect of concomitant energy deficit and exercise on skeletal muscle protein turnover and abundance.

In this study we provide an integrated physiological, endocrine and metabolic assessment in parallel with skeletal muscle dynamic proteome profiling in response to a 5-day pronounced energy deficit intervention with concomitant aerobic-type exercise, compared to an energy balance intervention in humans. Our analysis detected key physiological, endocrine and metabolic responses coherent with an energy preservation response triggered by energy deficit and increase in fatty acid oxidation, with skeletal muscle showing two distinct responses: a clear up-regulation of mitochondrial proteins synthesis and abundance, and a down-regulation of extracellular matrix and cytoskeletal proteins. Our findings corroborate and expand animal and human findings suggesting that energy deficit improves skeletal muscle quality and leads to a more youthful phenotype through enhancement of mitochondrial proteome quality and reduction extracellular matrix proteins associated to age-related fibrosis (Coen et al., 2015; Mahdy, 2019; Marosi et al., 2018; Rhoads et al., 2020).

These findings provide mechanistic support for the use of energy restrictive diets with concomitant exercise for the enhancement of skeletal muscle phenotype and yield insights into metabolic pathways with the potential to derive tailored therapeutic support to mimic the effects of energy deficit in skeletal muscle.

## RESULTS

Ten physically active, healthy men (participant characteristics are presented in supplementary Table 1) were studied during three consecutive 5-day periods that were designed to manipulate energy availability and exercise (Figure 1A, further detailed in *Protocol*). Participants were initially monitored during a lead-in period of free-living (FL; days −5 to 0) when their diet and exercise were not controlled. The subsequent experimental periods included controlled exercise under either energy balance (EB; day 0 to 5) or energy deficit (ED; days 5 to 10) conditions. During EB, energy intake matched the requirements for the maintenance of body-mass, whereas during ED each participant’s energy intake and energy availability were controlled at 35% and 22% of the EB period, respectively (Supplementary table 2).

**Figure 1.**
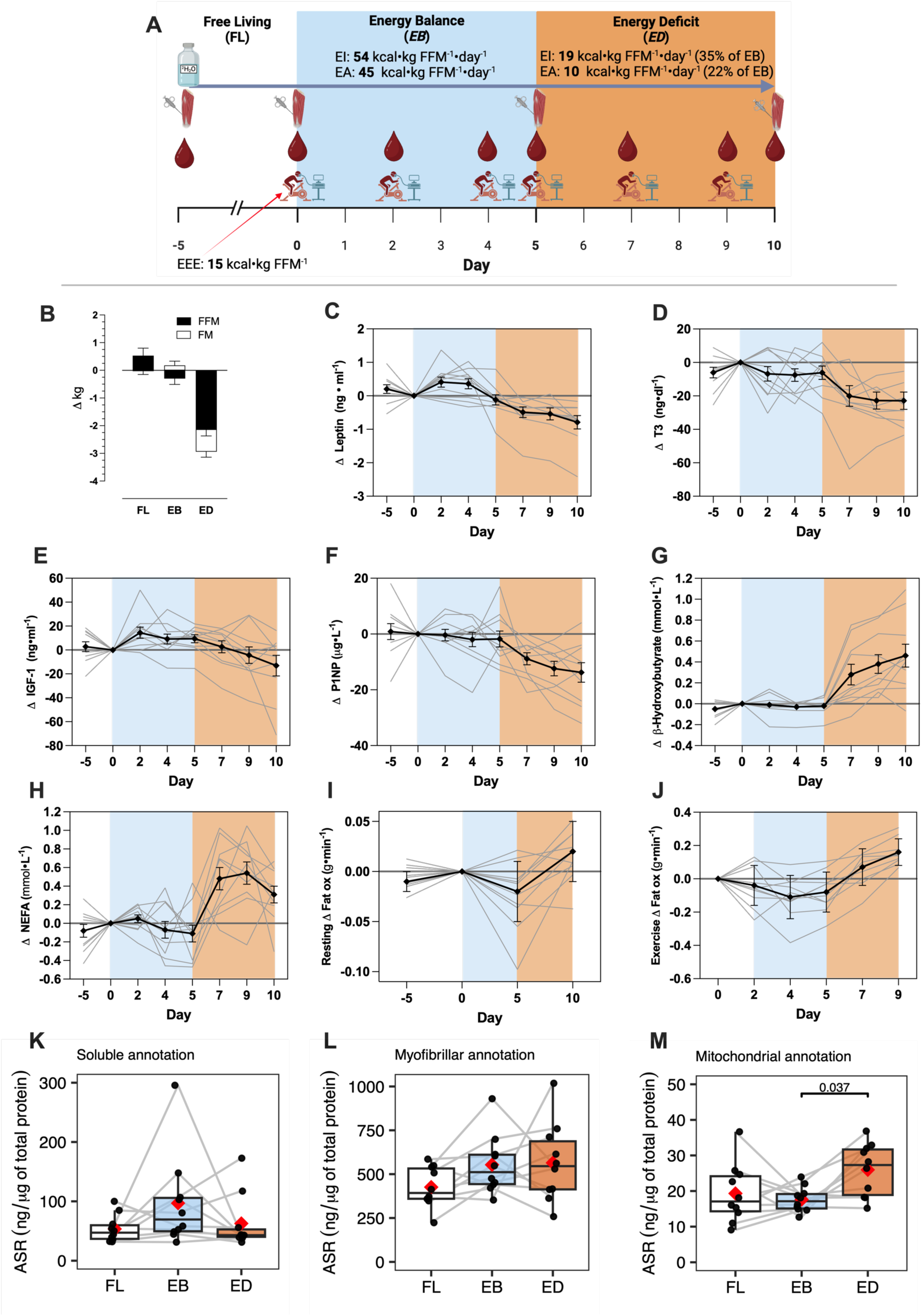
Study design and key metabolic variables summary. **A)** Schematic overview of the study design, created with BioRender.com. **B)** Change (mean ± SEM) of body fat free mass (FFM) and fat mass (FM) between the start and end of each 5-day period. **C – J:** Change (mean ± SEM) relative to Day 0 (beginning of controlled intervention phase) in **C)** plasma leptin, **D)** serum triiodothyronine (*T_3_*), **E)** serum insulin-like growth factor 1 (*IGF-1*), **F)** plasma procollagen type 1 N-terminal propeptide (*P1NP*), **G)** plasma β-Hydroxybutyrate, **H)** plasma non-esterified fatty acid (*NEFA*) concentrations, **I)** resting fat oxidation, and **J)** mean fat oxidation rates during steady state exercise at ∼60% VO_2max_ of ∼90 min exercise. Absolute values and statistical analyses are presented in supplementary table 3. n = 10 (individuals) x 4, 6 or 8 (time-points). Details of resting and exercise variables are presented in supplementary tables 3 and 4, respectively. **K – M:** Absolute synthetic rate (*ASR*) of proteins annotated into **K)** soluble, **L)** myofibrillar, and **M)** mitochondrial assessed *in silico*. The box plot represents the interquartile range (IQR; 25th–75th percentile), with the horizontal line indicating the median. Whiskers extend to the minimum and maximum values within 1.5× IQR. The red diamond represents the mean. Pairwise comparisons of the estimated marginal means for the experimental period were conducted using the Bonferroni adjustment for multiple comparisons in the linear mixed-effects model. EA, energy availability; EI, energy intake; EEE, exercise energy expenditure.

The participants’ body mass (average 77.3 kg, standard deviation s.d. 8.1 kg) did not change during the FL and EB periods, whereas participants lost an average 2.98 kg (s.d. 0.7 kg; P < 0.001) after ED (Figure 1B). Body mass loss after ED included reductions in fat free mass (2.1 ± 0.8 kg; P < 0.001), fat mass (0.8 ± 0.6 kg; P < 0.001), and total body water (1.2 ± 1 kg; P < 0.001; supplementary Table 3).

As per experimental design, during ED we observed the expected changes in endocrine and physiological blood markers associated with energy preservation in response to energy deficit, with metabolism switching from glucose to fat utilisation at rest and during exercise, resembling previous reports (Areta et al., 2021; Caldwell et al., 2024; Müller et al., 2015) (Figure 1, and Extended Data Figures 1 and 2; supplementary tables 3 and 4). Specifically, during energy deficit there was a decrease in leptin, triiodothyronine, insulin-like growth factor 1, and serum procollagen type I N-propeptide (Figure 1 C, D, E and F). The decrease in energy and macronutrient availability produced a metabolic shift in circulating substrate availability evidenced by an increase in circulating β-hydroxybutyrate concentrations and non-esterified fatty acids (Figure 1, G and H), and glycerol (Extended Data Figure 1, A) and a decrease in skeletal muscle glycogen content (Extended Data Figure 1, B), but no changes in glucose (Extended Figure 1, C) or insulin concentration (Extended Data Figure 1, D). Accordingly, there was an increase in fat oxidation after ED at rest (Figure 1, I, and also during exercise (Figure 1, J), mirrored by a decrease in carbohydrate oxidation at rest and during exercise (Extended Figure 2, A and B, respectively). We observed no changes in circulating testosterone, or EPO, a decrease in GDF-15 during EB and similar pattern of β-CTX increase during EB and ED (Extended Data Figure 1, E, F, G and H, respectively), and no changes in resting metabolic rate (Extended Data Figure 2, C and D).

**Figure 2.**
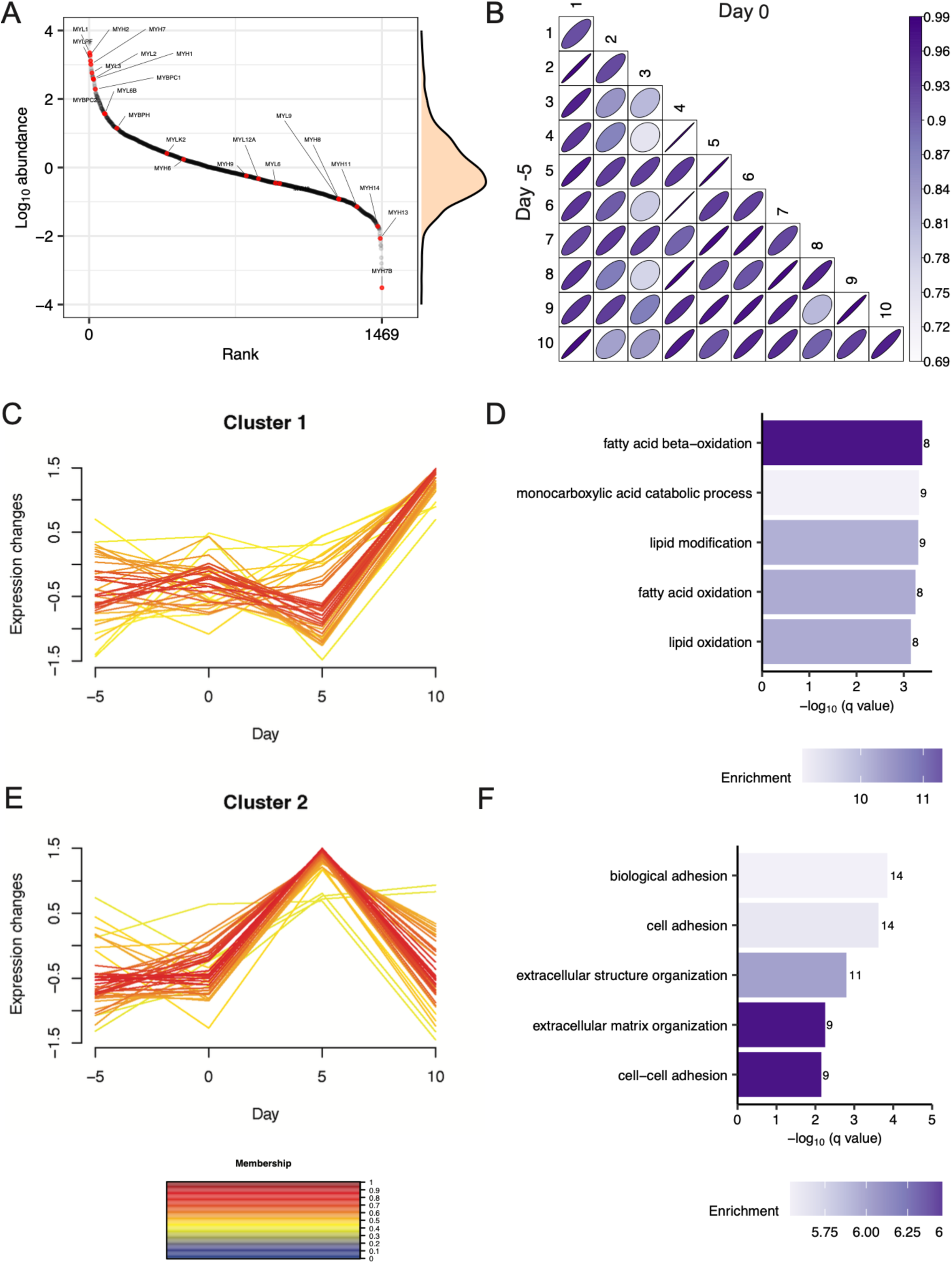
Unsupervised clustering analysis revealed two distinct time-dependent changes in skeletal muscle proteome abundance across a 5-day experimental period. **A)** rank distribution plot of mean protein abundance between subjects at Day −5 (n = 1469 proteins). Red data points highlight high abundance of myosin-related (muscle-specific) proteins. **B)** matrix correlation of abundance data between Day −5 and Day 0 within- and between-subjects, n = 1469 proteins, using Pearson’s correlation coefficient. C-means fuzzy clustering separated 108 statistically significant proteins (within subject one-way ANOVA P < 0.05) into 3 clusters with two major clusters; **C)** Cluster 1 (n=38) and **D)** Cluster 2 (n=44). Gene ontology (GO) analysis of Biological Process in proteins included in **E)** Cluster 1 and **F)** Cluster 2. GO terms were ranked by −log_10_ (q value) and the number of proteins included in each GO term reported alongside each entry. Each bar chart colour scale represents the level of GO enrichment.

### Energy deficit increases skeletal muscle mitochondrial protein synthesis

Across, FL, EB, and ED we measured absolute rates of protein synthesis of 736 proteins which were annotated in silico, and 388, 147 and 203 proteins were encompassed in soluble, myofibrillar, and mitochondrial annotation, respectively. There was no significant difference of average ASR in soluble (P= 0.17, Figure 1, K) and myofibrillar (P= 0.13, Figure 1, L) pools, but there was significant difference in the mitochondrial pool (P= 0.036, Figure 1,M) represented by significantly faster ASR in ED compared to EB (P=0.037). Detailed analysis of pathways regulated in protein ASR and abundance is shown in the following sections.

### Proteomic abundance and absolute synthesis rates show distinct responses to energy deficit

Proteomic analysis was conducted on 40 muscle samples taken at 4 time points (n=10 participants) prior to, between and after the 3 experimental periods (Figure 1A, and *Protocol*). Stringent filtering was used to ensure robust statistical analysis of the within-subject repeated measures experimental design. Overall, 1671 proteins were confidently (FDR <1 %) identified, and after excluding proteins that were not detected in each of the 4 serial samples 1469 proteins had complete within-subject data and were used for statistical analysis (Figure 2A). Protein abundance measurements were highly reproducible and the within-subject coefficient of variation (CV) across the FL condition (day −5 and day 0) was M = 15.3 % (inter-quartile range: 7 % to 30.8 %, Figure 2B).

Time-series analysis was conducted on protein abundance data and C-means fuzzy clustering was used to discover shared temporal patterns amongst 108 proteins that exhibited significant (within-subject ANOVA; P < 0.05, q< 0.41) differences across FL, EB and ED conditions. Unsupervised statistical methods, such as C-means fuzzy clustering, are blind to the biological role of proteins, yet functional annotation of the resulting clusters found strong biological networks with supporting muscle-specific literature on co-function amongst proteins that shared temporal relationships (Figure 2, D and F). Cluster 1 included proteins (n=38) associated with mitochondrial metabolism that were unchanged during the FL and EB periods but significantly increased in abundance after 5 days ED (Figure 2, C and D; Extended Data Figure 5). Cluster 2 included proteins (n=44) associated with the extracellular matrix, vesicles and exosomes that did not change during the FL period, increased during EB and decreased during ED (Figure 2, E & F; Extended Data Figure 6). Proteins (n=26) included in Cluster 3 exhibited heterogenous patterns of change, were less strongly clustered (had lower membership values) and did not include well-defined biological relationships (Extended Data Figure 7).

Overall, the majority (93%) of proteins were unaltered in abundance across FL, EB, and ED periods with highly reproducible proteome across the FL condition (day −5 and day 0). Coordinated changes to the protein abundance profile of human muscle during the ED period were compared against EB period.. There were 128 differences (P<0.05, q< 0.387) between EB and ED, including 84 proteins that were more abundant and 44 proteins that were less abundant in ED compared to EB (Figure 3A). GO terms that were significantly enriched amongst proteins that were more abundant in ED, included mitochondrion (GO:0005739, q=1.27E-08), fatty-acid beta-oxidation (GO:0006635, q=0.0037), carboxylic acid catabolic processes (GO:0046395, q=0.0038), monocarboxylic acid catabolic processes (GO:0072329, q=0.0041), small molecule metabolic processes (GO:0044281, q=0.0044) and organic acid catabolic process (GO:0016054, q=0.0048) (Figure 3A). Whereas, the GO term, extracellular structure organisation (GO:0043062, q=0.068), was enriched amongst proteins that were significantly less abundant in ED included (Figure 3A).

**Figure 3.**
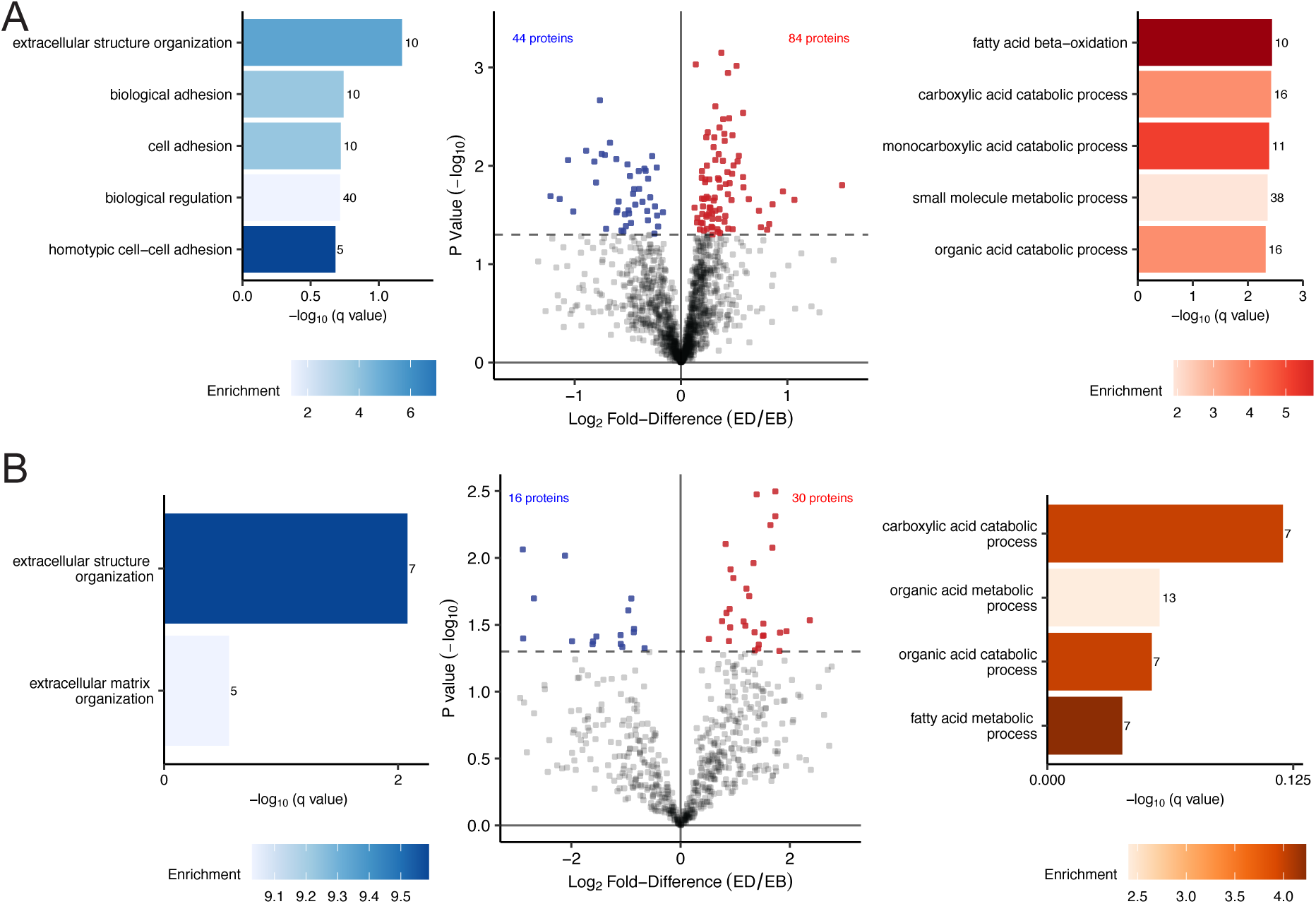
Energy deficit increased the abundance and absolute synthesis rates of proteins of mitochondrial metabolic processes and decreased the abundance and absolute synthesis rates of connective tissue. **A)** volcano plot comparing the Log_2_ Fold-Difference (ED/EB) protein abundance plotted against the −Log_10_ p value (n = 1469). Coloured data points represent proteins significantly more abundant (red, log_2_ Diff. > 0 and P < 0.05), less abundant (blue, log_2_ Diff. < 0 and P < 0.05), or stable (grey, P > 0.05) in ED compared to EB. **B)** volcano plot comparing the Log_2_ Fold-Difference (ED/EB) ASR plotted against the −Log_10_ p value (n = 625). Coloured data points represent proteins significantly faster ASR (red, log_2_ Diff. > 0 and P < 0.05), slower ASR (blue, log_2_ Diff. < 0 and P < 0.05), or stable (grey, P > 0.05) in ED compared to EB. The dashed horizontal line shows a threshold of statistical significance (P < 0.05). Gene ontology (GO) analysis of Biological Process in proteins significantly different between ween ED and EB. GO terms were ranked by−log_10_ (q value) and the number of proteins included in each GO term reported alongside each entry. Each bar chart colour scale represents the level of GO enrichment.

Protein-specific synthesis rates were investigated by biosynthetically labelling newly synthesised proteins using the stable isotope, deuterium oxide (D2O). Participants consumed D2O throughout the study period and a steady state enrichment of 0.84±0.11 % of the body water compartment was achieved. The absolute synthesis rate (ASR, ng/μg total protein) of individual muscle proteins was calculated from high-quality peptide mass isotopomer data collected from muscle samples that spanned each experimental condition. ASR data portrays both the rate of synthesis and abundance of each protein, which is essential to interpret differences in protein synthesis between muscles with different protein abundance profiles. Stringent filtering was used to ensure robust statistical analysis of 736 proteins that had complete protein synthesis data across FL, EB, and ED conditions.

To provide an overview, protein-specific ASR data were summed to reflect the bulk (mixed-protein) synthesis rate amongst myofibrillar, mitochondrial and sarcoplasmic muscle components. The human MitoCarta database was used to identify (n=203) mitochondrial proteins, the selection of myofibrillar (n=147) proteins was based on a manually curated subset of GO: 0030016 and the remaining (n=388) proteins were regarded as being sarcoplasmic. The sum ASR of mitochondrially annotated proteins was 47% greater (P=0.037) during ED (25.95 ± 7.42 ng/μg total protein /day) than EB (17.61 ± 3.51 ng/μg total protein/day), whereas there were no significant differences in the synthesis rate of mixed myofibrillar (515 ± 169 ng/μg total protein /day) or sarcoplasmic (71 ± 49 ng/μg total protein /day) proteins throughout the FL, EB and ED conditions (Figure 1, K, L, and M).

The more detailed effects of ED on skeletal muscle protein turnover were investigated by comparing protein-specific ASR data between EB and ED conditions. There were 46 differences (P<0.05, q< 0.30) between EB and ED, including 30 proteins with higher ASR and 16 proteins that with lower ASR in ED as compared to EB (Figure 3B). GO terms, including extracellular structure organization (GO:0043062, q=0.0083) and extracellular matrix organization (GO:0030198, q=0.28) were enriched amongst proteins with lower ASR during ED. Proteins that increased in ASR, included MYH7 and enzymes of fatty-acid beta-oxidation but GO terms were not statistically enriched amongst these proteins. MYH7 is a slow isoform of myosin heavy chain, expressed in slow-twitch oxidative muscle fibres, and the gain in both the abundance and synthesis rate of MYH7 during ED is consistent with studies on weight loss without exercise, reporting increases in muscle MYH7 mRNA expression alongside lower circulating levels of leptin and T3 (Baldwin et al., 2011; Rosenbaum et al., 2018).

### Energy deficit enhances muscle mitochondrial adaptations to exercise

Proteome profiling encompassed the majority of respiratory chain subunits and enzymes of the TCA cycle and fatty acid beta-oxidation which showed predominant increase in the abundance of proteins assessed (Figure 4 A, B, C and D). A common pattern of increases in the abundance and synthesis was evident across mitochondrial proteins specifically during ED (Figure 4E). The TCA cycle enzyme, citrate synthase (CS), is a commonly used biomarker of mitochondrial metabolism (Coen et al., 2015; Toledo et al., 2007) and was amongst the array of mitochondrial proteins that increased during ED (Figure 3 and Figure 4). Our findings agree with the effects of large-magnitude weight loss (7 to 21% body mass) achieved through energy deficit and exercise in people with obesity, which reported increases in mitochondrial respiratory capacity (Coen et al., 2015) and mitochondrial enzyme activity, including CS (Coen et al., 2015; Toledo et al., 2007). The mitochondrial adaptations to ED cooccurred with multilevel evidence of changes in lipid metabolism, including shifts increases in fat oxidation at rest and during exercise (Figure 1, I and J), circulating levels of non-esterified fatty acids (Figure 1, H) and ketones (Figure 1, G) and muscle perilipin (PLIN) proteins (Extended Figure 4), which are integral to the storage and breakdown of intramuscular triglyceride-containing lipid droplets (Barrett et al., 2022).

**Figure 4.**
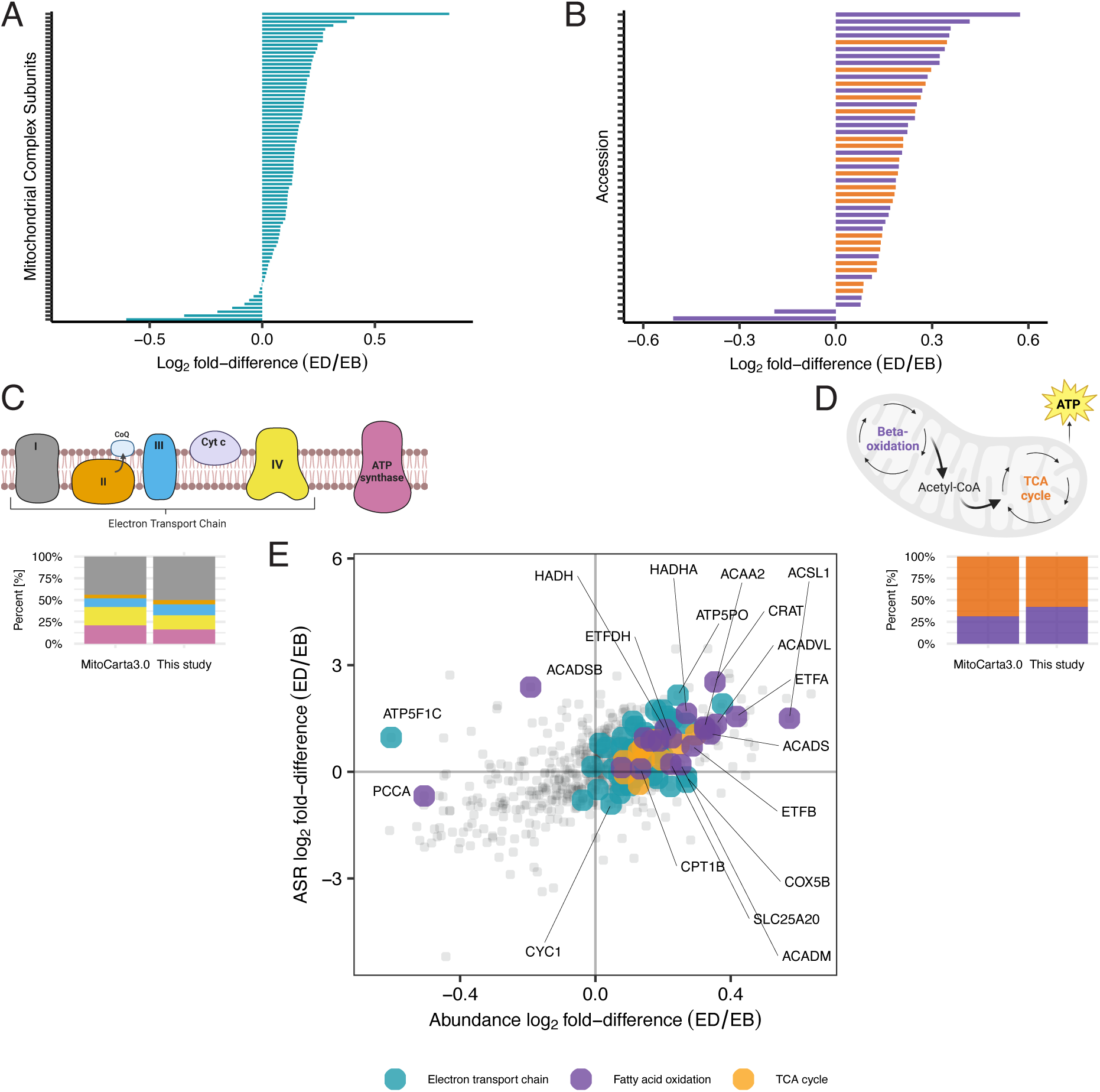
Energy deficit induces increase in abundance and ASR of proteins related to electron transport chain, fatty acid beta oxidation, and TCA cycle. Bar chart comparing the Log_2_ Fold-Difference (ED/EB) protein abundance of A, electron transport chains and B, fatty acid beta oxidation and TCA cycle. Stacked bar chat shows the relative distribution of mitochondrial proteins contained in and classified by the Human MitoCarta3.0 database for electron transport chain (C) and Beta-oxidation and TCA cycle (D). E, Scatter plot comparing the differences in the Log_2_ Fold-Difference (ED/EB) between protein abundance (x-axis) and protein ATR (y-axis). Proteins related to electron transport chain, fatty acid beta oxidation, and TCA cycle are coloured by blue, purple, and orange, respectively. Shiny app can be accessed via 10.6084/m9.figshare.27169632.

ED was associated with increases in the abundance of PLIN2 and PLIN5 and an increase in the ASR PLIN4. The discovery proteome abundance data were verified by orthogonal analysis using targeted mass spectrometry, which quantified PLIN2 and PLIN5 against stable isotope-labelled internal standards and verified both PLIN2 and PLIN5 significantly increased after ED (Extended Data Figure 4). Exercise training in humans increases the content of both PLIN2 and 5 proteins (Shepherd et al., 2013; Shepherd, Cocks, et al., 2017), which is likely to be an important adaptation to support greater intramuscular triglyceride storage, especially in the face of elevated free fatty acid availability produced by ED (Figure 1, H). Furthermore, PLIN5 is proposed to create a physical and metabolic link between lipid droplets and mitochondria, facilitating triglyceride hydrolysis and fatty acid oxidation (Wang et al., 2011).

PLIN4 is the most abundant perilipin in muscle but the role of PLIN4 in muscle metabolic responses to exercise is not yet clear. PLIN4 is primarily located to sarcolemma regions of slow twitch fibres (Pourteymour et al., 2015) and in adipocytes, PLIN4 coats nascent lipid droplets earlier than other PLIN proteins (Wolins et al., 2006). The muscle mRNA expression of PLIN4 decreases after 12 weeks of training (Pourteymour et al., 2015), whereas we (Shepherd, Strauss, et al., 2017) have reported a trend for greater PLIN4 protein abundance in the muscle of endurance-trained athletes. Our finding that PLIN4 synthesis increases without any change in PLIN4 abundance, together with the earlier observations, suggests the response of muscle PLIN4 to exercise involves a change in turnover rate rather than change in protein abundance.

Key proteins associated with mitochondrial quality, including LONP1, PITRM1, SSBP1, TIMM13, TIMM9, BDH1 and ALDH2 were enriched and had higher rates of synthesis during the ED compared to EB period. Elevations in BDH1, which is a component of muscle ketone body metabolism, link with the increased levels of circulating ketones (Figure 1G) and heightened fat utilisation at rest (Figure 1I) during exercise (Figure 1J) under the ED condition. In mice, ketone flux through BDH1 is essential to the adaptive response to intermittent time-restricted feeding (iTRF), and similar to our ED condition in humans, iTRF in mice increases muscle TCA cycle, OXPHOS and ETC protein abundance (Williams et al., 2024).

Muscle-specific knockout of BDH1 disrupts mitochondrial remodelling and is associated with lesser abundance of LONP1 (Williams et al., 2024), which agrees with our observations of parallel increases in BDH1 and LONP1 in human muscle during ED conditions. LONP1 is a mitochondrial protease involved in the activation of PINK1/Parkin pathway *(Zanini et al., 2023)* and, in mice, treadmill exercise increases LONP1 abundance alongside other markers of mitochondrial unfolded protein response (Cordeiro et al., 2020). High-intensity aerobic training in humans (Granata et al., 2021) did not change mitochondrial LONP1 abundance, which may point to a specific benefit of the ED intervention in humans.

The abundance of the mitochondrial matrix peptidase, PITRM1, also increased during ED and is positively associated with mitochondrial function. Humans with loss-of-function mutations of PITRM1 have markedly less active muscle respiratory chain complexes and, in mice, lower PITRM1 content is associated with lower levels of O_2_ utilisation (Brunetti et al., 2016). The benefits of exercise during ED extended to other mitochondrial protective mechanisms, including aldehyde dehydrogenase (ALDH2), which is an antioxidant enzyme that protects against lipid-peroxidation modifications to proteins. ALDH2 is more abundant in the muscle of high-capacity runner rats (Burniston et al., 2014), which are protected from cardiometabolic disease and muscle dysfunction. Moreover, overexpression of ALDH2 restores mitochondrial dysfunction, attenuates levels of muscle atrophy markers, and enhances physical capacity in mice (Q. Zhang et al., 2017). Proteins associated with mitochondrial DNA stability, including SSBP1 (Huang et al., 2009) and MSH5 (Bannwarth et al., 2012) also had higher levels of synthesis and abundance during ED and add evidence on the positive effects on mitochondria.

### The skeletal muscle extracellular matrix and cytoskeleton proteins are responsive to energy deficit

Proteins with roles in extracellular structure and organisation exhibited complex patterns of change that suggest reorganisation of the cytoskeleton and ECM during ED (Figure 3A, Figure 5, and Extended Data Figure 8). Decreases in the abundance and ASR of extracellular matrix-linked proteins (COL1A2, DCN, FN1, PRELP, S100A4, VTN, AZGP1, NID1, CLU, CLEC14, APOA1 and AGT, COL4A1, COL6A3 and LUM), and cytoskeletal proteins (ANXA2, EHBP1L1, FLNA, TBCB and VIM) cooccurred with increases in abundance and ASR of proteins associated with the dystrophin-associated protein complex (DPC) (DTNA, PLEC, LAMA2 and OBSCN) and tubulin polymerisation (CLIP1).

**Figure 5.**
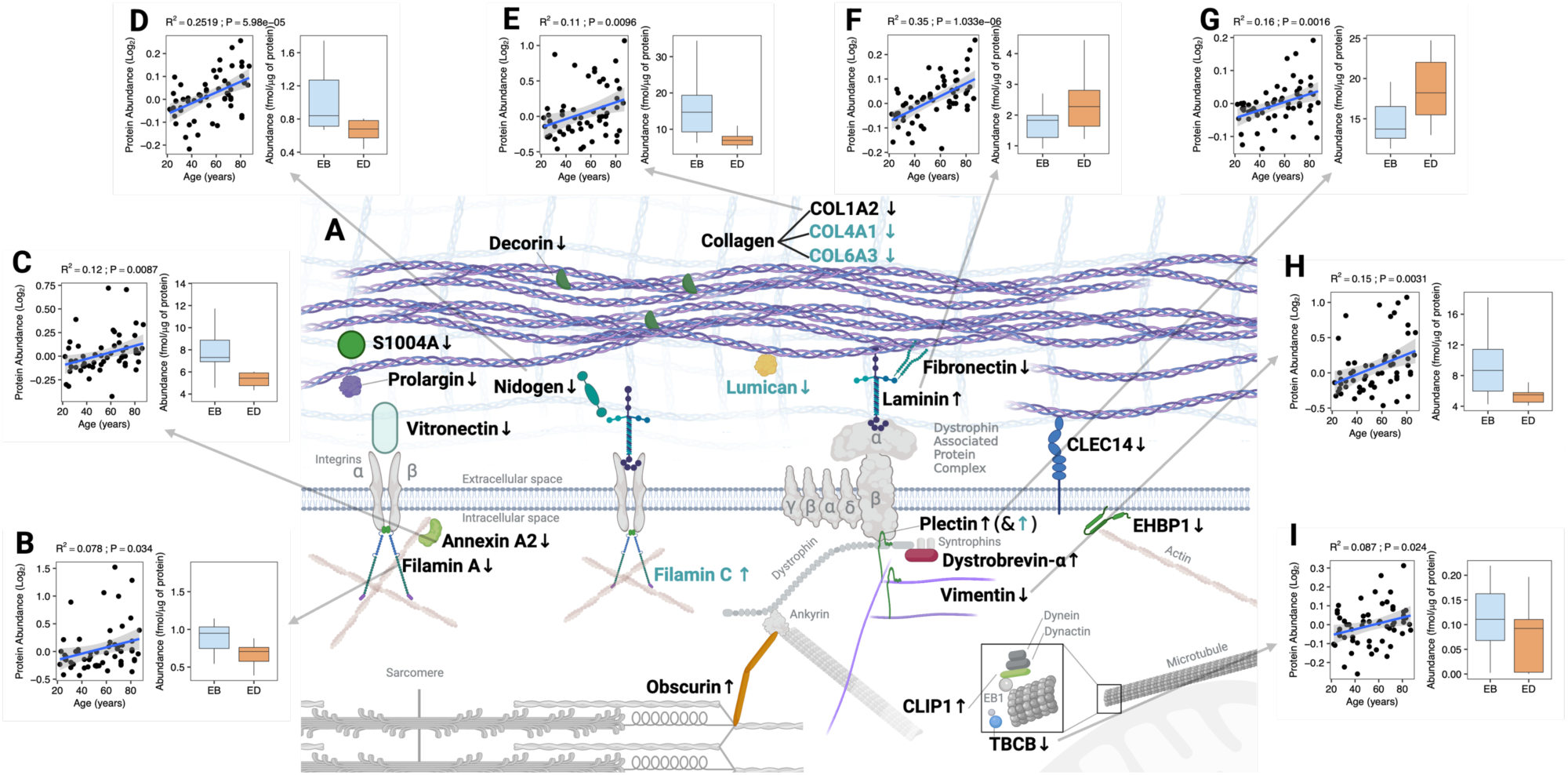
Schematic representation of extracellular matrix and cytoskeletal protein abundance (black text) and absolute synthesis rate (ASR; light emerald green text) changes in response to energy deficit (A), and relationship between abundance of proteins and ageing data generated by Ubaida-Mohien et al. (2019), compared to changes in abundance in our study (panels B to I). The majority of the proteins associated to the extracellular matrix (collagen —COL1A2, COL4A1 and COL6A3—, decorin, fibronectin, nidogen, vitronectin, prolargin, S100A4 and CLEC14A), as well as those of the intracellular space associated to the cytoskeleton (annexin, filamin A, EHBP1L1 and vimentin) were reduced in abundance or ASR during the energy deficit period. In contrast, proteins associated to the dystrophin associated protein complex (Laminin, dystrobrevin-α, plectin and obscurin) were increased. An exception is CLIP1, which may be related to an increase in stability of the polimerisation on the plus end of microtubules, and Filamin C. Proteins FLNA(B), ANXA2(C), NID1(D), COL1A2(E), LAMA2 (F), PLEC (G), VIM (H) and TBCB (F) and showed an abundance positively correlated with ageing (Ubaida-Mohien et al., 2019), a trend that was opposed by changes in abundance in COL1A2, NID1, FLNA, TBCB, VIM and ANXA2, but not LAMA2 or PLEC. Coloured proteins are proteins for which we report significant changes, light-grey/faded proteins are added for context.

Collagen is a major component of the extracellular matrix (Davis et al., 2013) and was particularly responsive to ED. Fibrillar collagen (COL1A2), network collagen (COL4A1, COL6A3) and proteins with roles in collagen binding and genesis (DCN, S100A4, PRLEP, NID1, FN1, VTN and CLEC14A, LUM) were less abundant or had lower rates of synthesis after ED. The decreases in collagen abundance and synthesis are consistent with the changes in circulating markers of collagen formation (P1NP, Figure 1F) and collagen breakdown (β-CTX, Extended Data Figure 1H) in ED. Furthermore, lumican decreased in ASR during ED and is a key extracellular matrix proteoglycan (Krishnan et al., 2012) and a myokine with effects on bone formation (Lee et al., 2020). The decrease in ASR of lumican cooccurred with decreases in abundance of ECM glycoproteins (decorin, vitronectin and nidogen), and mixed responses amongst basement membrane and cytoskeletal proteins. Laminin (LAMA2) is an important component of the basement membrane and increased in abundance, whereas the abundance of the basement membrane protein, prolargin, decreased. LAMA2 interacts with integrins and alpha-dystroglycan, and components of the DPC, including DTNA, OBSCN and PLEC also increased in abundance during ED. Other components of the DPC (SNTA1, SNTB1, DAG1 and DMD) were included in the proteomic data but did not change in abundance, and proteins with roles in the actin cytoskeleton (EHBP1L1, FLNA and ANXA2), intermediate filaments (VIM, FLNC) and microtubule network (CLIP1, TBCB) were also responsive to ED.

Ageing is associated with an accumulation of extracellular matrix proteins in muscle (Mahdy, 2019; McKiernan et al., 2012) that, in rhesus monkeys, can be countered by life-long caloric restriction (McKiernan et al., 2012; Rhoads et al., 2020). We investigated whether the muscle responses to ED reported here in young adults oppose the effects of ageing on extracellular matrix proteins across human adult ageing. Ubaida-Mohien et al. (2019) reports a significant positive relationship between ageing and the abundance of ECM proteins (FLNA, ANXA2, NID1, COL1A2, LAMA2, PLEC, VIM and TBCB) in skeletal muscle. We found 6 (FLNA, ANXA2, NID1, COL1A2, VIM and TBCB) out of 8 of these proteins exhibited the opposite effect (decreased in abundance) after ED (Figure 5B-F). These contrasting effects of ageing and energy deficient on the human muscle proteome add support to the experimental observations on life-long energy deficit in rhesus monkeys (McKiernan et al., 2012; Rhoads et al., 2020).

## DISCUSSION

Skeletal muscle is central to metabolic regulation and locomotion, yet the impact of acute energy deficit on muscle phenotype in exercising humans remains poorly defined. We show that a 5-day period of pronounced energy restriction (78% reduction in energy availability), combined with aerobic-type exercise, remodels the skeletal muscle proteome in young healthy adults. This remodelling involves a shift toward a more oxidative phenotype, with increased synthesis of mitochondrial proteins and elevated abundance of enzymes involved in fatty acid β-oxidation, the tricarboxylic acid cycle, electron transport, and mitochondrial quality control, alongside enhanced glycogen and lipid storage. Notably, these adaptations occurred without reductions in bulk myofibrillar or sarcoplasmic protein synthesis. To the best of our knowledge, we are also the first to report decreased abundance of extracellular matrix proteins and increased levels of components of the dystrophin-associated protein complex in human skeletal muscle following energy deficit. These coordinated proteomic responses suggest an improvement in muscle quality and metabolic efficiency, with potential implications for interventions aimed at enhancing healthspan, lifespan, and physical capacity.

Our data offers new molecular insights into the mitochondrial adaptations associated with energy deficit in exercising humans. Consistent with studies involving substantial weight loss in overweight or obese individuals showing increases in the content and activity of some mitochondrial enzymes (Coen et al., 2015; Toledo et al., 2008), we observed a coordinated increase in mitochondrial protein synthesis resulting in upregulation of a broad array of mitochondrial proteins. Notably, 287 out of 343 quantified mitochondrial proteins were increased in abundance, reflecting widespread remodelling of the mitochondrial proteome. Prominent protein networks regulated by energy deficit were associated with an enhancement of mitochondrial proteostasis, mtDNA-quality control and protein import, which agrees with transcriptomic responses reported in humans following 2 years of 12% caloric restriction with no exercise (Das et al., 2023). A possible key candidate we identify mediating this response is BDH1, which has been linked to increases in fatty acid oxidation efficiency and exercise capacity with intermittent-time-restricted feeding-induced energy deficit in mice (Williams et al., 2024). These findings support a synergistic role of energy restriction and exercise in promoting mitochondrial biogenesis and are aligned with murine data showing increased mitochondrial content and improved physical capacity (Marosi et al., 2018). Improvements in mitochondrial metabolism by energy deficit have also been linked to the maintenance of a youthful skeletal muscle phenotype in life-long energy restricted macaque monkeys, reflected in a reduction in the age-related decline muscle contractile area resulting from fibrosis (Rhoads et al., 2020), which is aligned with our observations of the extracellular matrix and cytoskeletal protein response.

Our results also point to notable changes in ECM and cytoskeletal proteins, which may be relevant for muscle aging and fibrosis. Skeletal muscle fibrosis is a hallmark of aging (Evans & Lexell, 1995; Mahdy, 2019), and lifelong caloric restriction has been shown to prevent fibrotic accumulation in monkeys (McKiernan et al., 2012; Rhoads et al., 2020). Here, we present the first human data indicating that short-term energy deficit directly reduces ECM proteins associated with aging, including COL1A2 and nidogen, as well as others such as decorin, S100A4, prolargin, vitronectin, fibronectin, and CLEC14. In contrast, proteins associated with the dystrophin-associated protein complex (e.g., laminin, plectin, dystrobrevin-alpha, and obscurin) increased in abundance, suggesting enhanced membrane stability and cell adhesion in response to ECM remodeling. These findings raise the possibility that energy deficit could therapeutically reduce pathological ECM accumulation linked to impaired glucose metabolism and sarcopenia (Draicchio et al., 2022; Melouane et al., 2020).

The modulation of ECM proteins during ED aligns with recent plasma proteome data from individuals undergoing prolonged fasting, where extracellular proteins were enriched (Pietzner et al., 2024). The downregulation of non-essential structural proteins under energy deficit, combined with endocrine adaptations such as reduced T3, leptin, and IGF-1, and enhanced fat oxidation and aerobic metabolic pathway activity, aligns with trade-offs predicted by life-history theory (Pontzer & McGrosky, 2022), whereby there may be a reallocation of energetic resources for processes that are immediately important for survival. In this context, our findings suggest that during energy-limited states, skeletal muscle prioritizes the maintenance of contractile and metabolic capacity—particularly when stimulated by exercise. This interpretation of the findings provides a conceptual framework to further enquire about the potential functional and health effects of GLP-1 agonist-induced weight-loss on skeletal muscle in the context of the current debate (Conte et al., 2024; Prado et al., 2024). Our findings highlight the importance of looking beyond reductions in bulk protein synthesis and muscle size with energy deficit and for future research to focus on select metabolic, non-metabolic and functional changes of skeletal muscle to understand the potential positive and negative effects of energy deficit on muscle and overall health.

Our study has limitations. The quasi-experimental design restricts causal inference, and the small sample size (n=10), consisting solely of healthy young males, limits generalizability. While participants maintained perceived effort during exercise, we did not formally assess physical performance. The intervention’s short duration and severity provided a useful model to detect molecular changes, but future research should examine longer or clinically relevant protocols. Although macronutrient intake was standardized as a percentage of total energy intake, varying protein levels could have affected protein synthesis (Areta et al., 2014).

In conclusion, our findings reveal a dynamic skeletal muscle proteome response to short-term energy deficit combined with exercise, characterized by enhanced mitochondrial remodelling and reductions in extracellular matrix proteins. These adaptations suggest a shift toward improved metabolic health and resistance to muscle aging, even in the face of hormonal and metabolic signals favouring energy conservation. Such responses likely reflect a conserved evolutionary strategy for maintaining physical capacity under energetic constraint. As energy-restricted states become increasingly relevant due to lifestyle, medical, and pharmacological interventions, understanding their effects on muscle health is critical. Our work offers a molecular foundation for future studies aimed at leveraging energy deficit to optimize skeletal muscle function and overall health.

## METHODS

### Participants

Ten healthy males that were active (completing >3 sessions/hours of aerobic exercise per week), weight-stable (reported ≤ 2 kg fluctuation, over the previous 6 months) without known health complications were recruited for the study (see supplementary table 1) after being pre-screened and providing a written consent. Ethical approval for the study was granted by the NHS North-West-Liverpool Central Research Ethics Committee (Ref: 21/NW/0205) and the study was registered on the clinical trials register (ClinicalTrials.gov ID: NCT05203133). Participant characteristics are presented in supplementary Table 1, and Figure 1 A provides a schematic of the experiment design (further detailed in *Protocol*).

### Study Design

After participant characterisation testing, participants followed a quasi-experimental design intervention (**Figure 1, A**) during a period of 15 consecutive days, which consisted of three consecutive 5-day periods of, 1) Free living (Days −5 to −1), followed by two phases of controlled diet and exercise 2) Energy Balance (*EB*: Days 0 to 5), which aimed at maintaining body mass, and 3) Energy deficit (ED: Days 5 to 10), which induced weight-loss through a dietary energy deficit (Supplementary table 2). We sampled fasting blood, resting metabolic rate, and respiratory gases during morning exercise, and skeletal muscle (vastus lateralis of quadriceps) before and after each period (Figure 1 A; and Protocol).

### Participant characterisation assessment

Prior to the start of the study, participants reported to the laboratory for assessment of anthropometric parameters, cycloergometer-based (Lode Corival cpet: Lode, Groningen, Netherlands) maximal oxygen consumption (VO_2Max_; Moxus Modular Metabolic System, AEI Technologies, Pittsburgh, PA, USA), peak power output (PPO), lactate threshold (LT; Biosen C-Line, EKF Diagnostic, Cardiff, UK) and submaximal O_2_ utilisation using standard protocols to determine participant fitness and the workload for exercise intensity during intervention (Newell et al., 2015).

### Dietary and Exercise Intervention phase with manipulation of EA (days 0 to 10)

From day 0 to day 10, all food was provided to participants in custom-made, pre-packaged meals, and participants consumed all and only the foodstuff provided, with non-calorie containing drinks allowed *ad libitum*. This period consisted of two 5-day phases; days 0 to 4 (inclusive) comprised the Energy Balance (*EB*) dietary provision, with days 5 – 9 (inclusive), of Energy Deficit (ED) diet containing 35% of the energy intake of EB. Proportion of macronutrients was clamped at 60% carbohydrate, 20% fat and 20% protein in both diets (see diet details in Supplementary Table 2). During each phase, individuals completed three cycling sessions to expend 15 kcal·kg Fat Free Mass^−1^·day^−1^. Average daily energy availability ((Energy intake – Exercise Energy Expenditure)·kg FFM^−1^) (Loucks et al., 1998) achieved was 45 and 10 kcal·kg FFM^−1^·day^−1^ for the EB & ED phases, respectively, which represents a 78% reduction in energy availability in ED.

### Study Intervention Phase (days −5 to 10): assessments and biological samples collection

For all testing sessions (days −5, 0, 2, 4, 5, 7, 9 and 10), participants arrived at the laboratory at 0700-0800 hrs in a fasted state. Resting metabolic rate (RMR) was assessed using indirect calorimetry (GEM Open Circuit Indirect Calorimeter; GEMNutrition Ltd., Warrington, UK) following a standard 25-minute protocol (Iraki et al., 2021). Body composition was assessed via bio-electrical impedance analysis (*BIA*; SECA mBCA 515: SECA GMBH, Hamburg, Germany), on each laboratory visit, and via a whole-body dual-energy X-ray absorptiometry scan (*DXA*; Hologic, Manchester, United Kingdom) on days −5, 0, 5, and 10 using standardized procedures (Nana et al., 2016). Fasting venous blood samples were collected at each laboratory visit, and skeletal muscle was obtained from the lateral portion of the vastus lateralis of the quadriceps muscle after each RMR assessment, and prior to exercise.

#### Exercise protocol

Exercise sessions were initiated with consecutive three-minute stages at 50 W, 75 W, 100 W and 125 W. Thereafter workload was set to a steady-state power output to elicit 60% VO_2Max_ until achieving a target EEE of 15 kcal·kg FFM^−1^·day^−1^. Respiratory gas assessments were repeated every 15 min with rate of perceived exertion (RPE (Borg, 1982)), assessed at the end of each stage. Fat and carbohydrate oxidation were estimated based on the equations standard equations (Jeukendrup & Wallis, 2005). Participants were provided with a carbohydrate-rich meal (30 g carbohydrates, <1 g fat and 3 g protein) at ∼5-15 min before exercise and at 5 min breaks after 30 and 60 min of exercise.

### Blood analyses

Analysis of blood metabolites, hormones and bone turnover markers: Plasma glucose, lactate, non-esterified fatty acids (NEFA) and glycerol concentrations were analysed using commercially available kits and a Randox Daytona spectrophotometer (Randox, Crumlin, Ireland). Commercially available enzyme-linked immunosorbent assays (ELISA) were used to measure serum total triiodothyronine (T3; DiaMetra S.r.l, Perugia, Italy), serum insulin-like growth factor 1 (IGF-1), plasma insulin, and plasma leptin (all DRG International Inc., Springfield, NJ, USA), serum erythropoietin (EPO; Abcam, Cambridge, UK) and serum growth/differentiation factor-15 (GDF-15; R&D Systems, Minneapolis, MN, USA). Bone turnover markers (plasma ß-CTX and P1NP), and serum total testosterone concentrations were analysed by a commercial laboratory (Liverpool Clinical Laboratories, Liverpool, UK).

### Skeletal Muscle Proteomic Analysis

Stable isotope labelling in vivo. Biosynthetic labelling of newly synthesised proteins was achieved by oral consumption of deuterium oxide (D_2_O; Sigma-Aldrich, UK). A ‘loading’ + ‘maintenance’ design was used to rapidly raise the enrichment of D_2_O in the participants body water compartment and then sustain a steady-state level of enrichment throughout the experimental period. After collection of the baseline muscle sample on experimental day −5, participants consumed doses of 200 ml, 150 ml and 150 ml of 99.8 atom % of D2O at approximately 3-hour intervals across the day. On subsequent days, participants were provided with 50 ml ‘maintenance doses’ of 99.8 atom % of D2O in sterile feeding bottles and were prompted to consume one dose each day by members of the research team.

Calculation of D2O Enrichment. Body water enrichment of D2O was measured in plasma samples against external standards that were constructed by adding D2O to 18 MΩ water over the range from 0.0 to 5.0 % in 0.5 % increments. D2O enrichment of aqueous solutions was determined by gas chromatography-mass spectrometry after exchange with acetone (McCabe et al., 2006). Samples were centrifuged at 12,000 g, 4°C for 5 min, and 20 µl of plasma supernatant or standard was reacted overnight at room temperature with 2 µl of 10 M NaOH and 4 µl of 5% (v/v) acetone in acetonitrile. Acetone was then extracted into 500 µl chloroform and water was captured in 0.5 g Na2SO4 before transferring a 200 µl aliquot of chloroform to an auto-sampler vial. Samples and standards were analysed in triplicate by using an Agilent 5973 N mass selective detector coupled to an Agilent 6890 gas chromatography system (Agilent Technologies, Santa Clara, CA, USA). A CD624-GC column (30 m 30.25 mm 31.40 mm) was used in all analyses. Samples (1 ul) were injected by using an Agilent 7683 auto sampler. The temperature program began at 50°C and increased by 30°C/min to 150°C and was held for 1min. The split ratio was 50:1 with a helium flow of 1.5 ml/min. Acetone eluted at 3 min. The mass spectrometer was operated in the electron impact mode (70 eV) and selective ion monitoring of m/z 58 and 59 was performed by using a dwell time of 10 ms/ ion.

Protein Extraction and Quantification. Proteins were extracted from muscle samples as previously described (1, 2). Muscle samples were ground in liquid nitrogen, then homogenized on ice in 10 volumes of 1 % Triton X-100, 50 mM Tris, pH 7.4 (including complete protease inhibitor; Roche Diagnostics, Lewes, United Kingdom) using a PolyTron homogenizer (KINEMATICA, PT 1200 E) followed by sonication. Homogenates were incubated on ice for 15 min, then centrifuged at 1000 x g, 4 °C, for 5 min to fractionate myofibrillar (pellet) from soluble (supernatant) proteins. Myofibrillar proteins were resuspended in a half-volume of homogenization buffer followed by centrifuged at 1000 x g, 4 °C, for 5 min. The washed myofibrillar pellet was then solubilized in lysis buffer (7 M urea, 2 M thiourea, 4% CHAPS, 30 mM Tris, pH 8.5). Aliquots of protein were precipitated in 5 volumes of ice-cold acetone and incubated for 1 h at −20 °C, and resuspended in 200 uL of lysis buffer. Proteins were cleared by centrifugation at 12,000 x g, 4 °C, for 45 min. Total protein concentration (μg/μl) was quantified against bovine serum albumin (BSA) standards using the Bradford assay (Thermo Scientific, #23236), according to the manufacturer’s instructions.

Protein Digestion. Tryptic digestion was performed using the filter-aided sample preparation (FASP) method (3). Aliquots containing 100 µg protein were washed with 200 µl of UA buffer (8 M urea, 100 mM Tris, pH 8.5). Proteins were incubated at 37 °C for 15 min in UA buffer containing 100 mM dithiothreitol followed by incubation (20 min at 4 °C) protected from light in UA buffer containing 50 mM iodoacetamide. UA buffer was exchanged for 50 mM ammonium bicarbonate and sequencing-grade trypsin (Promega, Madison, WI, USA) was added at an enzyme to protein ratio of 1:50. Digestion was allowed to proceed at 37 °C overnight then peptides were collected in 100 μl of 50 mM ammonium bicarbonate containing 0.2 % (v/v) trifluoroacetic acid. Samples containing 4 µg of peptides were de-salted using C18 Zip-tips (Millipore Billercia, MA, USA) and eluted in 40% (v/v) acetonitrile and 0.1 % (v/v) trifluoroacetic acid. Peptides were dried by vacuum centrifugation and resuspended in 20 µl of 2.5 % (v/v) acetonitrile, 0.1 % (v/v) formic acid containing 10 fmol/ μl yeast alcohol dehydrogenase (MassPrep standard; Waters Corp., Milford, MA).

Liquid Chromatography-Mass Spectrometry Analysis. Peptide mixtures were analysed using an Ultimate 3000 RSLC nano liquid chromatography system (Thermo Scientific) coupled to Q-Exactive orbitrap mass spectrometer (Thermo Scientific). Samples were loaded on to the trapping column (Thermo Scientific, PepMapTM 100, 5 μm C18, 300 μm X 5 mm), using ulPickUp injection, for 1 minute at a flow rate of 25 μl/min with 0.1 % (v/v) TFA and 2% (v/v) ACN. Samples were resolved on a 500 mm analytical column (Easy-Spray C18 75 μm, 2 μm column) using a gradient of 97.5 % A (0.1 % formic acid) 2.5 % B (79.9 % ACN, 20 % water, 0.1 % formic acid) to 50 % A 50 % B over 150 min at a flow rate of 300 nl/min. The data-dependent selection of the top-10 precursors selected from a mass range of m/z 300-1600 was used for data acquisition consisted of a 70,000-resolution full-scan MS scan at m/z 200 (AGC set to 3e6 ions with a maximum fill time of 240 ms). MS/MS data were acquired using quadrupole ion selection with a 3.0 m/z window, HCD fragmentation with a normalized collision energy of 30 and in the orbitrap analyzer at 17,500-resolution at m/z 200 (AGC target 5e4 ion with a maximum fill time of 80 ms). To avoid repeated selection of peptides for MS/MS, the program used a 30 s dynamic exclusion window.

Label-Free Quantitation of Protein Abundances. Progenesis Quantitative Informatics for Proteomics (QI-P; Nonlinear Dynamics, Waters Corp., Newcastle, UK, Version 4.2) was used for label-free quantitation, consistent with previous studies. Log-transformed MS data were normalized by inter-sample abundance ratio, and relative protein abundances were calculated using nonconflicting peptides only. In addition, abundance data were normalized to the 3 most abundant peptides of yeast ADH1 to obtain abundance estimates in fmol/μg total protein (4). MS/MS spectra were exported in Mascot generic format and searched against the Swiss-Prot database (2021_03) restricted to Homo-sapiens (20,371 sequences) using locally implemented Mascot server (v.2.2.03; www.matrixscience.com). The enzyme specificity was trypsin with 2 allowed missed cleavages, carbamidomethylation of cysteine (fixed modification) and oxidation of methionine (variable modification). M/Z error tolerances of 10 ppm for peptide ions and 20 ppm for fragment ion spectra were used. The Mascot output (xml format), restricted to non-homologous protein identifications was recombined with MS profile data in Progenesis.

### Parallel reaction monitoring

For targeted analysis of PLIN2 and PLIN5 in human skeletal muscle, the instrument was operated in parallel reaction monitoring (PRM) mode using an Ultimate 3000 RSLCTM nano system (Thermo Scientific) coupled to a Q-Exactive orbitrap mass spectrometer (Thermo Scientific). Stable isotopically-labeled (SIL) peptides corresponding to the targeted endogenous peptides were spiked in each sample as a 1 fmol/uL concentration and 0.5 fmol/uL for PLIN2 and PLIN5, respectively (HeavyPeptideTM AQUA Basic). Samples (4.5 μL corresponding to 1.8 ng of protein) were loaded on to the trapping column (Thermo Scientific, PepMap100, 5 μm C18, 300 μm X 5 mm), using μlPickUp injection, for 1 minute at a flow rate of 100 μL/min with 2 % (v/v) acetonitrile and 0.1 % (v/v) formic acid. Samples were resolved on a 500 mm analytical column (Easy-Spray C18 75 μm, 2 μm column) using a gradient of 97.5 % A (0.1 % formic acid) 2.5 % B (79.9 % ACN, 20 % water, 0.1 % formic acid) to 55 % A 45 % B over 31 min at a flow rate of 300 nL/min. Untargeted MS/MS analysis of peptides were acquired via HCD fragmentation mode (normalised collision energy 27) and detection in the Orbitrap (70,000 resolution at m/z 200, 5e4 automatic gain control (AGC), 240 ms maximum injection time, and 2 m/z isolation window) and a 30 s dynamic exclusion window was used to avoid repeated selection of peptides for MS/MS analysis.

Data were processed using Skyline-daily v23.1.1.353. Product ion extraction ion chromatograms (XICs) were generated by including all matching scans for MS2 filtering. Ion match tolerance was set to 0.055 m/z and matched to 1+ for MS2 filtering of b- or y-type ions that were reliably quantitative in all samples. All data were manually confirmed for co-elution of MS1 and MS2 and ratio dot product (rdotp) ≥ 0.91 in all samples. Quantification was performed with MS2 fragments only and absolute abundance (fmol) was estimated by comparing the ratio of the signal intensities between the light peptide (endogenous) and heavy SIL peptide (L/H ratio) using a one-point calibration.

Measurement of protein turnover rates. Mass isotopomer abundance data were extracted from MS spectra using Progenesis Quantitative Informatics (Non-Linear Dynamics, Newcastle, UK). Consistent with previous work {Camera, 2017 #959;Hesketh, 2020 #1509;Nishimura, 2023 #10}, the abundances of peptide mass isotopomers were collected over the entire chromatographic peak for each proteotypic peptide that was used for label-free quantitation of protein abundances. Mass isotopomer information was processed using in-house scripts written in Python (version 3.12.4). The incorporation of deuterium into newly synthesised protein was assessed by measuring the increase in the relative isotopomer abundance (RIA) of the m_1_ mass isotopomer relative to the sum of the m_0_ and m_1_ mass isotopomers (Equation 3) that exhibits rise-to-plateau kinetics of an exponential regression {Sadygov, 2020, Partial Isotope Profiles Are Sufficient for Protein Turnover Analysis Using Closed-Form Equations of Mass Isotopomer Dynamics} as a consequence of biosynthetic labelling of proteins *in vivo*.

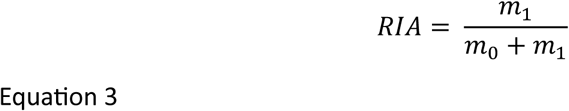

The plateau in RIA (*RIA_plateau_*) of each peptide was derived (Equation 4) from the total number (*N*) of ^2^H exchangeable H—C bonds in each peptide, which was referenced from standard tables {Holmes, 2015 #1044}, and the difference in the D:H ratio (^2^H/^1^H) between the natural environment (*DH_nat_*) and the experimental environment (*DH_exp_*) based on the molar percent enrichment of deuterium in the precursor pool, according to {Ilchenko, 2019, Calculation of the Protein Turnover Rate Using the Number of Incorporated 2H Atoms and Proteomics Analysis of a Single Labelled Sample}.

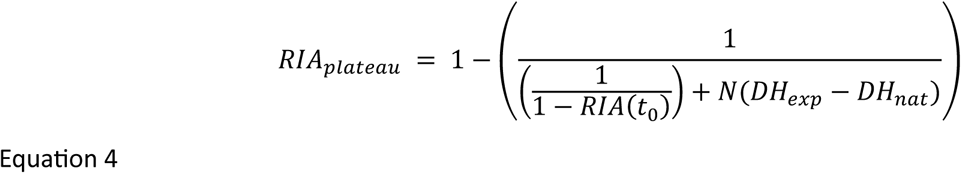

The rate constant of protein degradation (*k_deg_)* was calculated (Equation 5) between the beginning (t_0_) and end (t_1_) of each 5-day labelling period. Calculations for exponential regression (rise-to-plateau) kinetics reported in {Ilchenko, 2019, Calculation of the Protein Turnover Rate Using the Number of Incorporated 2H Atoms and Proteomics Analysis of a Single Labelled Sample} were used and *Kdeg* data were adjusted for differences in protein abundance (P) between the beginning (t_0_) and end (t_1_) of each labelling period.

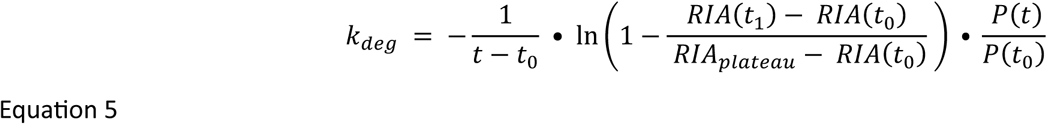

The absolute abundance (P) of each protein was calculated from the relative abundance (fmol/ μg total protein) measured by LFQ multiplied by the molecular weight (MW; kDa) of the protein (referenced from the UniProt Knowledgebase). Absolute synthesis rates (ASR) were derived (Equation 6) by multiplying peptide *K_deg_* by the absolute abundance of the protein at the end of the labelling period P(t).

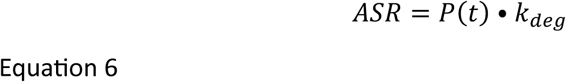

### Statistical Analyses

With the exception of skeletal muscle proteome analysis, data were collated using Microsoft Excel. Statistical analyses were conducted using IBM SPSS Statistics (v. 29.0.0.0, IBM, Armonk, NY, USA) and figures were created using Prism (v. 10.4.1, Graphpad Software Inc., San Diego, California, USA). Anthropometric, resting metabolic rate, exercise indirect calorimetry data and changes in all blood-borne markers were explored via linear mixed models. Significance was set at P < .05 for all statistical tests and data are reported as means ± SDs.

All statistical analysis of proteomic data was performed in R version 4.3.2 (2023-10-31). Within-subject one-way ANOVA was used to assess protein abundance and ASR. Differences in the abundance or ASR of proteins between EB and ED were reported as log_2_ transformed data and statistical significance was set at P < 0.05. A false discovery rate was not set, instead, q values (Storey & Tibshirani, 2003) at the P = 0.05 were reported. Pairwise comparisons of the estimated marginal means for the experimental period were conducted using the Bonferroni adjustment for multiple comparisons in the linear mixed-effects model with the emmeans R package (Lenth, 2025).

### Bioinformatic Analysis

Skeletal muscle proteomic data was processed and analysed using R (Version 4.2.2) and figures were generated using the R package, ggplot2 (Wickham & Sievert, 2009). The Search Tool for the Retrieval of INteracting Genes/proteins (STRING, Version 11.5) (6) was used to investigate protein interaction networks and calculate the enrichment of gene ontologies or KEGG pathways. Gene ontology analysis (GO) was performed via Overrepresentation Enrichment Analysis(B. Zhang et al., 2005) using Gene Ontology enRIchment anaLysis and visuaLizAtion tool (Gorilla) (Eden et al., 2009) corrected against the experiment-specific background consisting of all proteins that were included in statistical analysis. Enrichment of GO terms was considered significant if the Benjamin Hochberg adjusted p value was 0.01. Protein interaction networks were visualized by the Cytoscape string app (Doncheva et al., 2019) using the Omics Visualizer app (Legeay et al., 2020). The Mfuzz R package (Kumar & Futschik, 2007) was used to perform soft clustering analysis using the fuzzy c-means clustering algorithm. The minimum membership value for inclusion into a cluster was set at 0.25. The coverage of mitochondrial Complex subunit proteins, fatty acid beta oxidation, and TCA cycle was survey as identified in Human MitoCarta 3.0 (Rath et al., 2021).

## Supporting information

Protocol

## DATA AVAILABILITY

The mass spectrometry proteomics data generated in this study have been deposited to the ProteomeXchange Consortium via the PRIDE (Perez-Riverol et al., 2019) partner repository with the dataset identifier PXD061267 and 10.6019/PXD061267. Raw files of parallel reaction monitoring, Skyline document, processed results used as input for figure generation are available on PanoramaWeb (Sharma et al., 2014) (https://panoramaweb.org/margulis2025.url) and ProteomeXchange with the identifier PXD061311. R script and raw data used to generate the Shiny app were deposited in figshare (DOI: 10.6084/m9.figshare.27169632).

## AUTHOR CONTRIBUTIONS

Conceptualization: J.L.A. Data curation/software: Y.N., C.L.E.,H.L.T., J.G.B. and J.L.A. Formal analysis: Y.N., C.L.E., H.L.T., J.G.B. and J.L.A. Methodology: Y.N.,H.L.T.,W.L.F., J.S., S.S., J.G.B. and J.L.A. Visualization: Y.N. and J.L.A. Funding acquisition: J.G.B. and J.L.A. Project administration: J.L.A. Supervision: C.L.E., J.P.M., J.G.B. and J.L.A. Writing original draft: Y.N., J.G.B. and J.L.A. Writing review and editing: Y.N.,C.L.E, H.L.T.,W.L.F., J.P.M, J.S., S.S., J.G.B. and J.L.A.

## ACKNOWLEDGMENTS

We thank the participants for their dedication and effort in the participation of this project. This research was supported by data generated by Intramural Research Program of the NIH, National Institute on Aging. We thank Prof. Luigi Ferrucci for providing access to the skeletal muscle proteomics ageing database, which enabled the secondary analysis of human muscle ageing data in this manuscript. This work was supported by the Alliance for Potato Research and Education, and the Research Institute for Sport and Exercise Sciences at Liverpool John Moores University.

## Extended Data

**Extended Data Figure 1.**
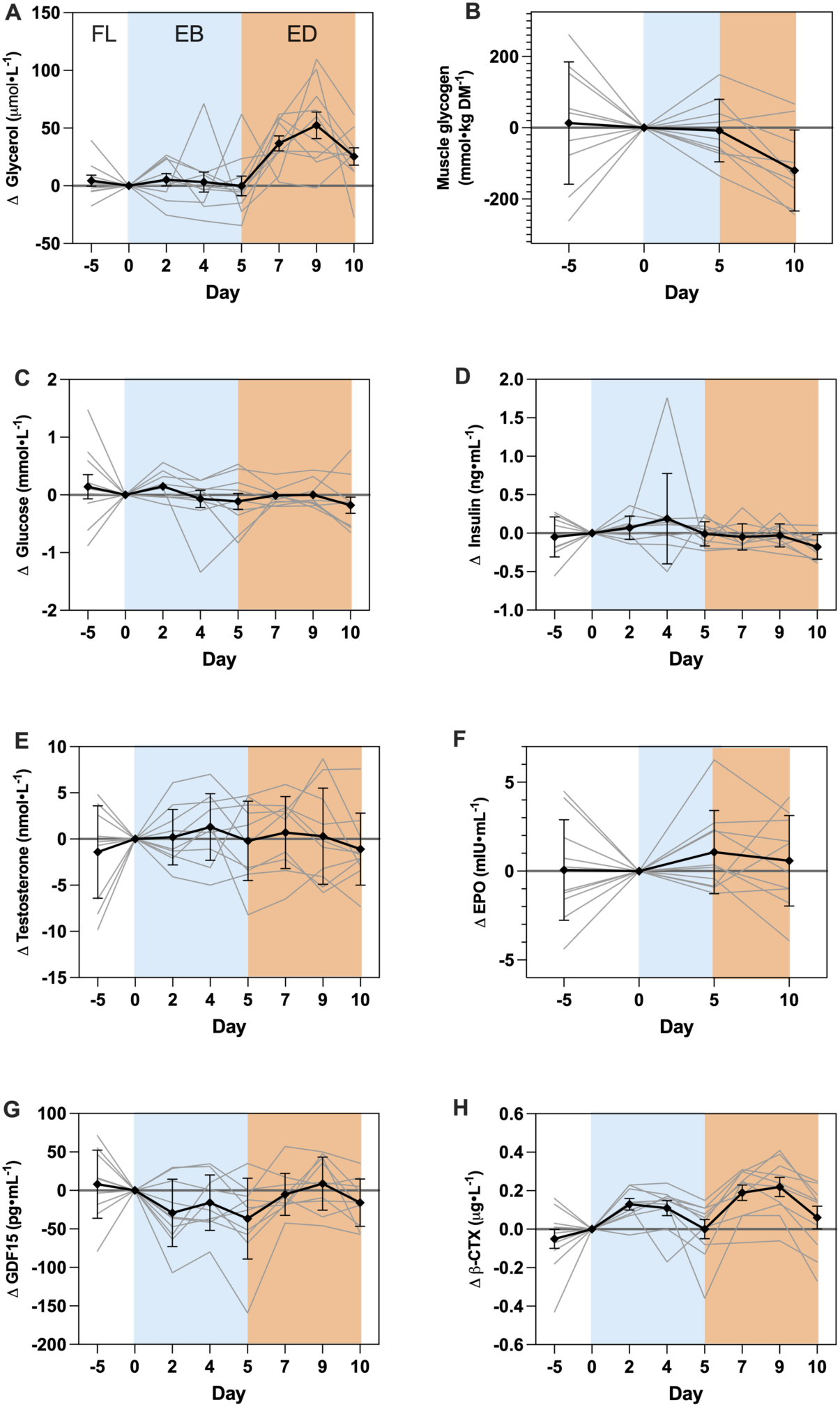
Blood metabolites and hormones and skeletal muscle glycogen. Change in **A)** plasma glycerol, **B)** Skeletal muscle glycogen, **C)** plasma glucose, **D)** plasma insulin, **E)** serum testosterone, **F)** serum erythropoietin, **G)** serum GDF-15 and **H)** plasma β-CTX. Absolute values and statistical analyses are presented in supplementary table 3. Details of exercise variables are presented in supplementary table 4. n = 10 (individuals) x 4 or 8 (samples).

**Extended Data Figure 2.**
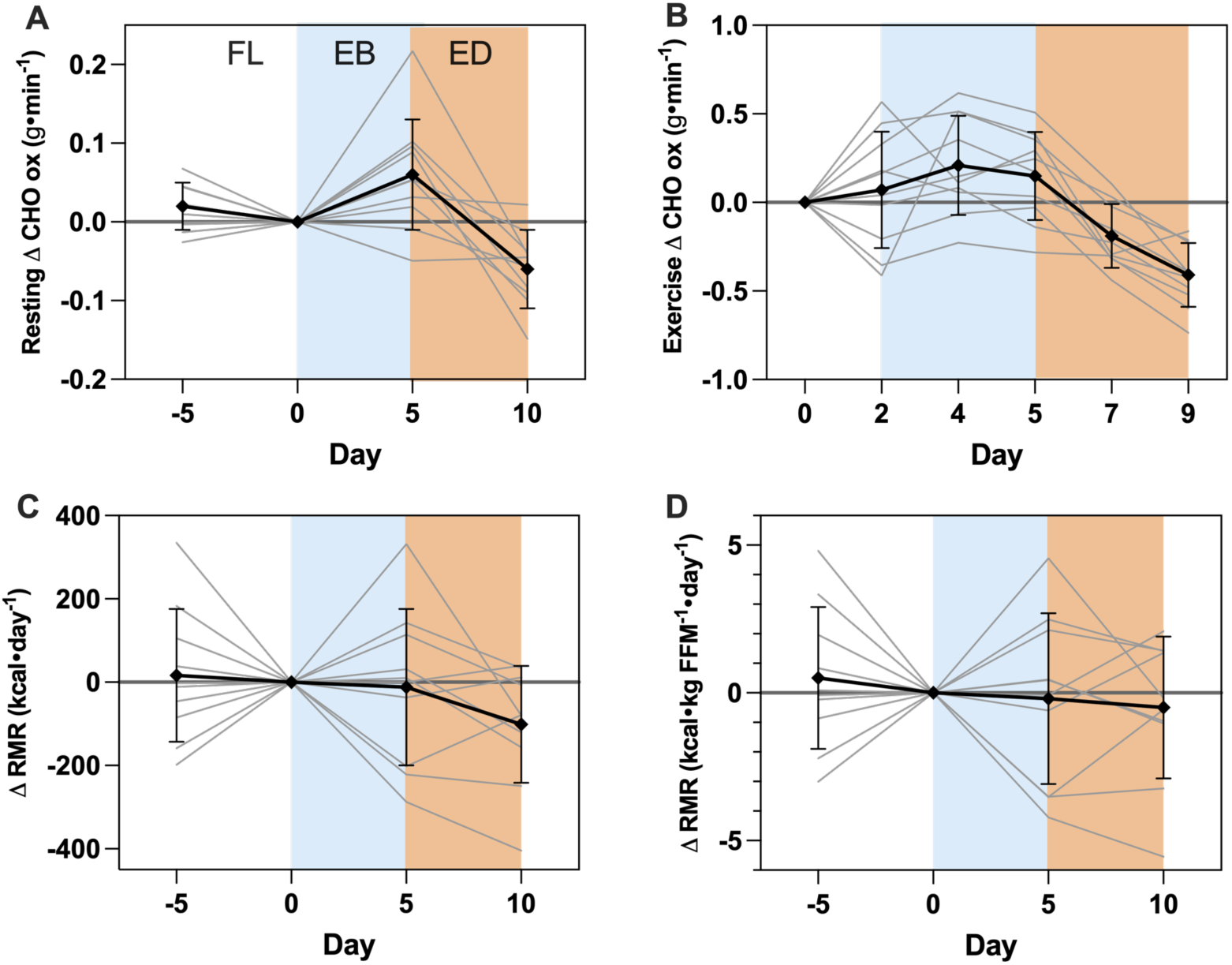
Resting and exercise-related respiratory parameters. Change in **A)** Resting carbohydrate (CHO) oxidation at rest, **B)** CHO oxidation during exercise, **C)** Resting metabolic rate and **D)** Resting metabolic rate (relative to fat free mass). Absolute values and statistical analyses are presented in supplementary table 3. Details of exercise variables are presented in supplementary table 4. n = 10 (individuals) x 4 or 6 (samples)

**Extended Data Figure 3.**
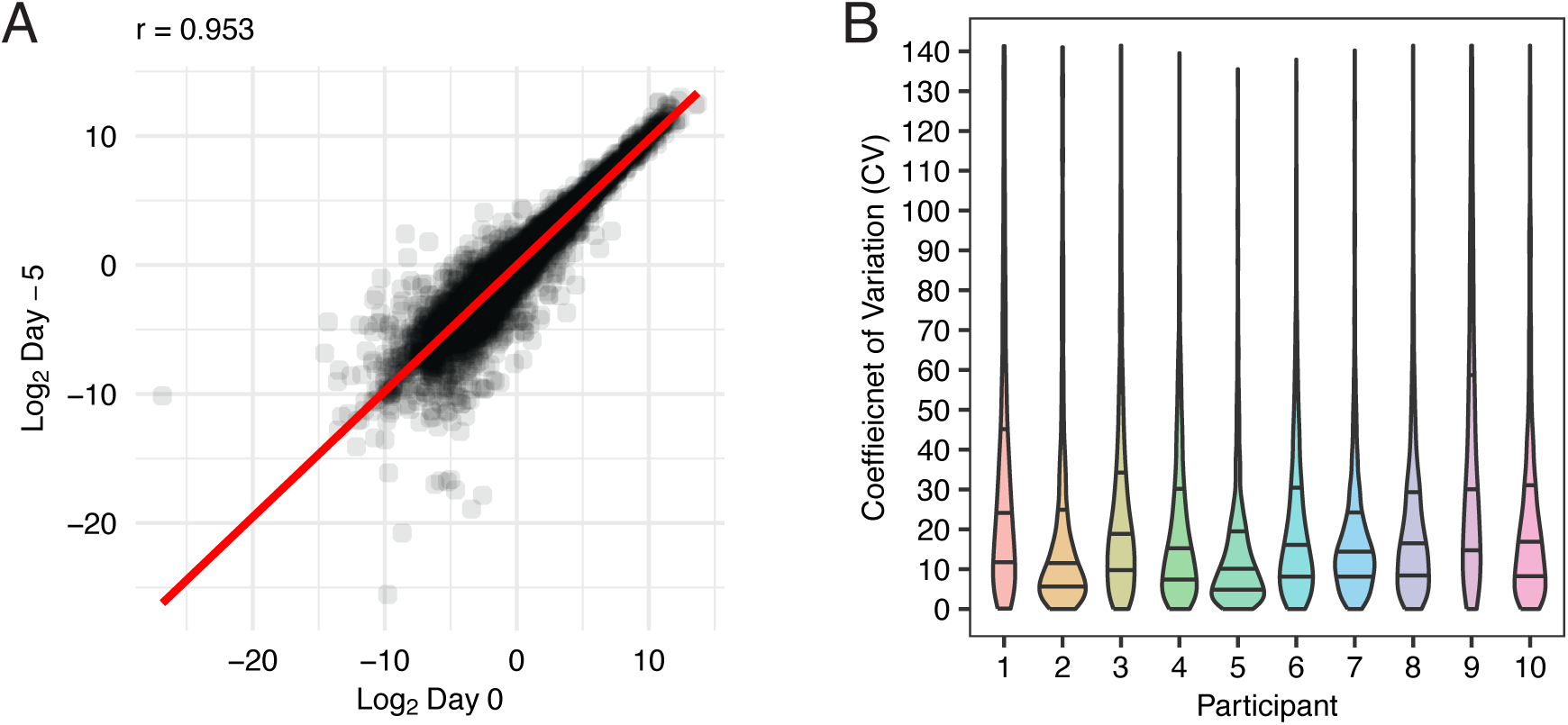
Skeletal muscle proteome abundance is stable during the FL condition. A, a scatter plot of protein abundance data between Day −5 and day 0 from all 10 participants. B, violin plots of Coefficient of Variation between Day −5 and Day 0 on a protein-by-protein basis.

**Extended Data Figure 4.**
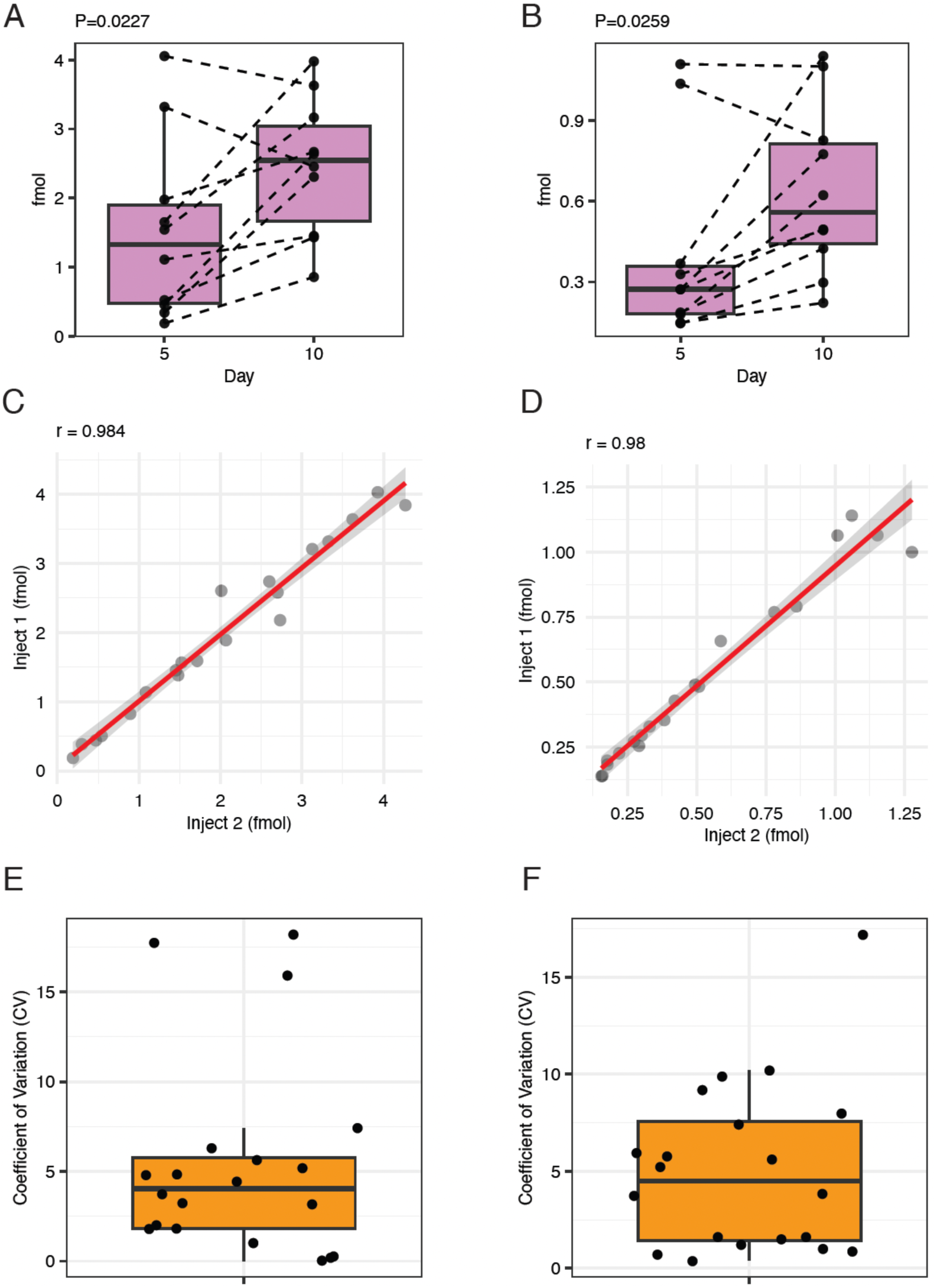
Targeted proteomic analysis via parallel reaction monitoring verifies PLIN2 and PLIN5 are more abundant in energy deficit. Box plot illustrating absolute protein abundance of A, PLIN2 and B, PLIN5. A scatter plot comparing absolute protein abundance between two independent injections into LC-MS in C, PLIN2 and D, PLIN5. Coefficient of Variation (CV) of absolute protein abundance between 2 independent injections into LC-MS in E, PLIN2 and F, PLIN5.

**Extended Data Figure 5.**
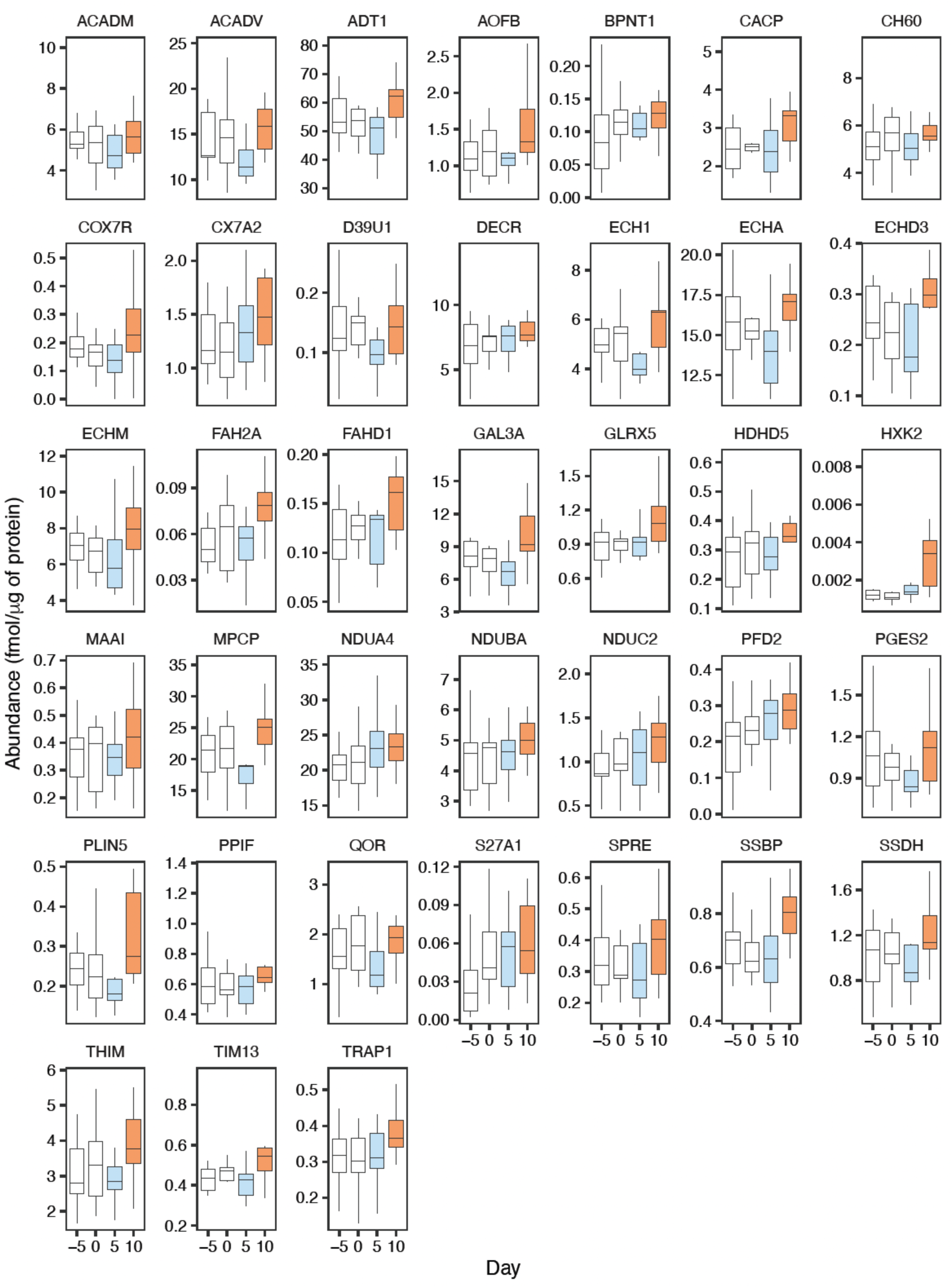
Box plots of proteins in cluster 1 identified by c-means fuzzy clustering. Box plots illustrating changes of protein abundance across the experimental period (38 proteins, all P<0.05). The box plot represents the interquartile range (IQR; 25th–75th percentile), with the horizontal line indicating the median. Whiskers extend to the minimum and maximum values within 1.5× IQR.

**Extended Data Figure 6.**
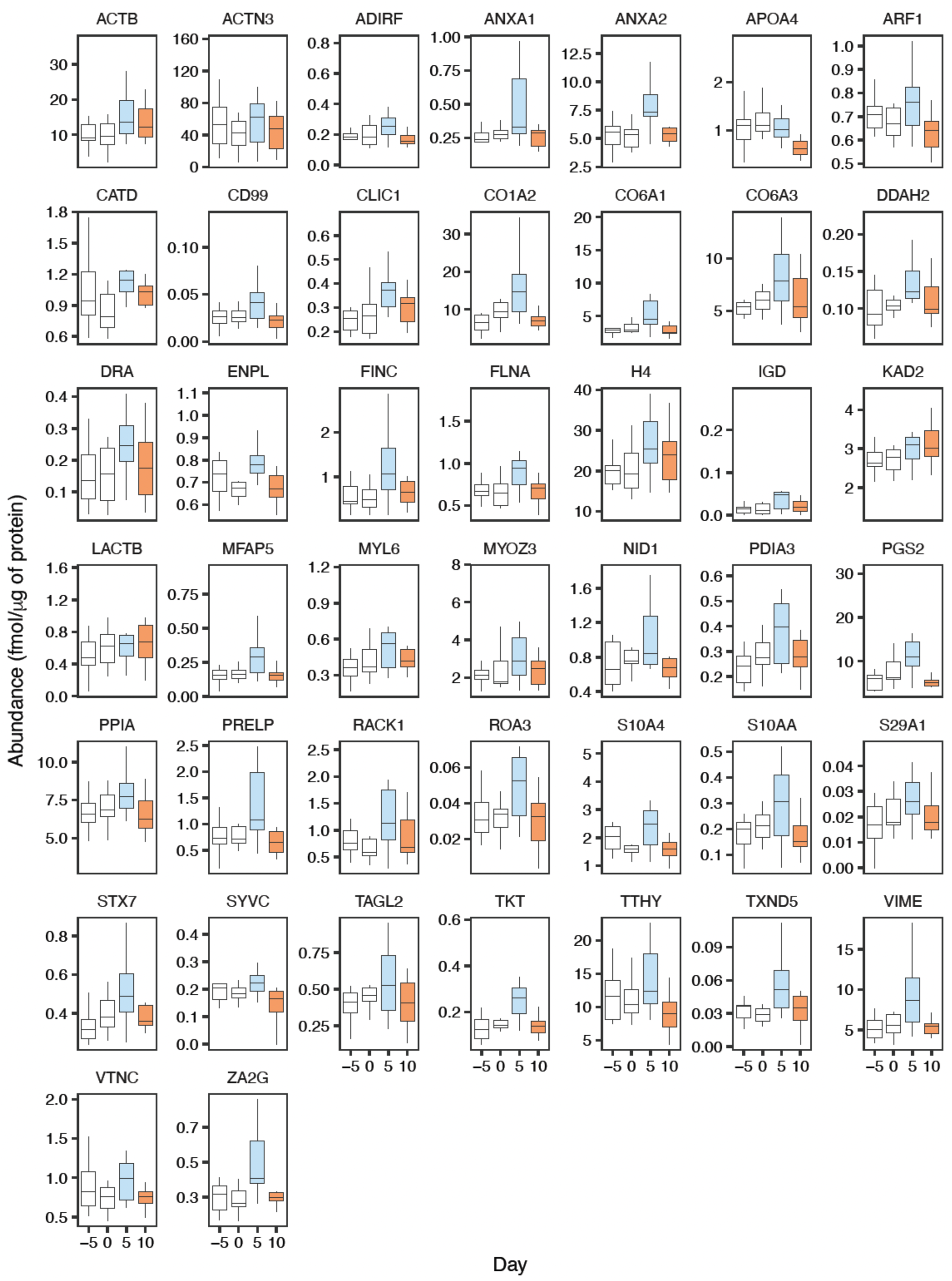
Box plots of proteins in cluster 2 identified by c-means fuzzy clustering. Box plots illustrating changes of protein abundance across the experimental period (44 proteins, all P<0.05). The box plot represents the interquartile range (IQR; 25th–75th percentile), with the horizontal line indicating the median. Whiskers extend to the minimum and maximum values within 1.5× IQR.

**Extended Data Figure 7.**
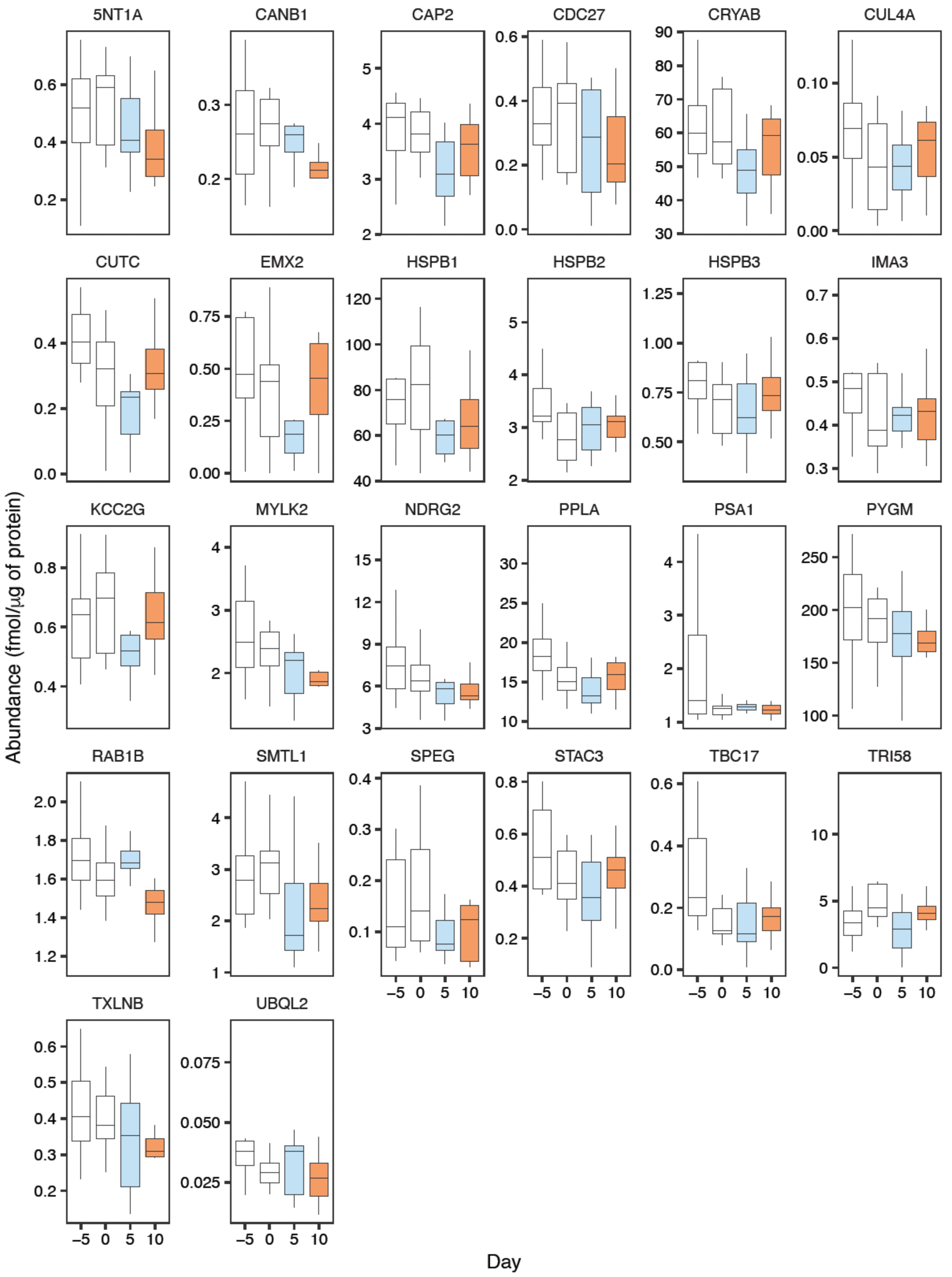
Box plots of proteins in cluster 3 identified by c-means fuzzy clustering. Box plots illustrating changes of protein abundance across the experimental period (26 proteins, all P<0.05). The box plot represents the interquartile range (IQR; 25th–75th percentile), with the horizontal line indicating the median. Whiskers extend to the minimum and maximum values within 1.5× IQR.

**Extended Data Figure 8.**
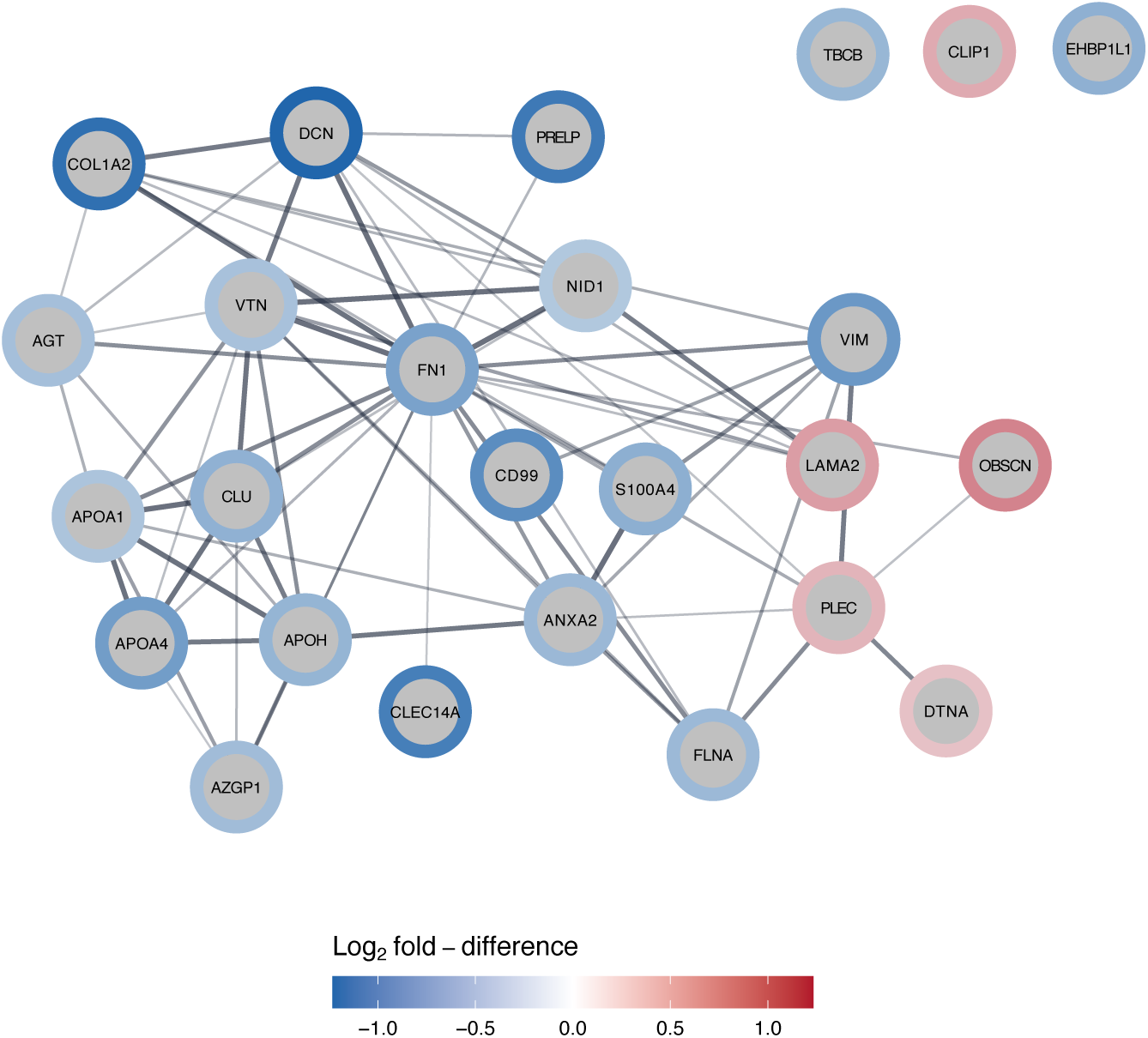
STRING network of proteins belonging to the extracellular matrix and cytoskeletal network identified through GO network analyses and independent analysis of significantly different proteins between EB and ED (Supplementary table 6). The boarder colour represents Log_2_ fold-difference in protein abundance between ED and EB.

## REFERENCES

Areta, J. L., Burke, L. M., Camera, D. M., West, D. W. D., Crawshay, S., Moore, D. R., Stellingwerff, T., Phillips, S. M., Hawley, J. A., & Coffey, V. G. (2014). Reduced resting skeletal muscle protein synthesis is rescued by resistance exercise and protein ingestion following short-term energy deficit. American Journal of Physiology-Endocrinology and Metabolism, 306(8), E989–E997. 10.1152/ajpendo.00590.2013

Areta, J. L., Taylor, H. L., & Koehler, K. (2021). Low energy availability: History, definition and evidence of its endocrine, metabolic and physiological effects in prospective studies in females and males. European Journal of Applied Physiology, 121(1), 1–21. 10.1007/s00421-020-04516-0

Baldwin, K. M., Joanisse, D. R., Haddad, F., Goldsmith, R. L., Gallagher, D., Pavlovich, K. H., Shamoon, E. L., Leibel, R. L., & Rosenbaum, M. (2011). Effects of weight loss and leptin on skeletal muscle in human subjects. American Journal of Physiology-Regulatory, Integrative and Comparative Physiology, 301(5), R1259–R1266. 10.1152/ajpregu.00397.2011

Bannwarth, S., Figueroa, A., Fragaki, K., Destroismaisons, L., Lacas-Gervais, S., Lespinasse, F., Vandenbos, F., Pradelli, L. A., Ricci, J.-E., Rötig, A., Michiels, J.-F., Vande Velde, C., & Paquis-Flucklinger, V. (2012). The human MSH5 (MutS Homolog 5) protein localizes to mitochondria and protects the mitochondrial genome from oxidative damage. Mitochondrion, 12(6), 654–665. 10.1016/j.mito.2012.07.111

Barrett, J. S., Whytock, K. L., Strauss, J. A., Wagenmakers, A. J. M., & Shepherd, S. O. (2022). High intramuscular triglyceride turnover rates and the link to insulin sensitivity: Influence of obesity, type 2 diabetes and physical activity. Applied Physiology, Nutrition, and Metabolism, 47(4), 343–356. 10.1139/apnm-2021-0631

Beals, J. W., Kayser, B. D., Smith, G. I., Schweitzer, G. G., Kirbach, K., Kearney, M. L., Yoshino, J., Rahman, G., Knight, R., Patterson, B. W., & Klein, S. (2023). Dietary weight loss-induced improvements in metabolic function are enhanced by exercise in people with obesity and prediabetes. Nature Metabolism, 5(7), 1221–1235. 10.1038/s42255-023-00829-4

Bergouignan, A. (2016). Towards human exploration of space: The THESEUS review series on nutrition and metabolism research priorities. Npj Microgravity.

Borg, G. A. (1982). Psychophysical bases of perceived exertion. Med Sci Sports Exerc, 14(5), 377–381.

Bramble, D. M., & Lieberman, D. E. (2004). Endurance running and the evolution of Homo. Nature, 432(7015), 345–352. 10.1038/nature03052

Brunetti, D., Torsvik, J., Dallabona, C., Teixeira, P., Sztromwasser, P., Fernandez-Vizarra, E., Cerutti, R., Reyes, A., Preziuso, C., D’Amati, G., Baruffini, E., Goffrini, P., Viscomi, C., Ferrero, I., Boman, H., Telstad, W., Johansson, S., Glaser, E., Knappskog, P. M., … Bindoff, L. A. (2016). Defective PITRM 1 mitochondrial peptidase is associated with Aβ amyloidotic neurodegeneration. EMBO Molecular Medicine, 8(3), 176–190. 10.15252/emmm.201505894

Burniston, J. G. (2019). Investigating Muscle Protein Turnover on a Protein-by-Protein Basis Using Dynamic Proteome Profiling. In J. G. Burniston & Y.-W. Chen (Eds.), Omics Approaches to Understanding Muscle Biology (pp. 171–190). Springer US. 10.1007/978-1-4939-9802-9_9

Burniston, J. G., Kenyani, J., Gray, D., Guadagnin, E., Jarman, I. H., Cobley, J. N., Cuthbertson, D. J., Chen, Y.-W., Wastling, J. M., Lisboa, P. J., Koch, L. G., & Britton, S. L. (2014). Conditional independence mapping of DIGE data reveals PDIA3 protein species as key nodes associated with muscle aerobic capacity. Journal of Proteomics, 106, 230–245. 10.1016/j.jprot.2014.04.015

Caldwell, H. G., Jeppesen, J. S., Lossius, L. O., Atti, J. P., Durrer, C. G., Oxfeldt, M., Melin, A. K., Hansen, M., Bangsbo, J., Gliemann, L., & Hellsten, Y. (2024). The whole-body and skeletal muscle metabolic response to 14 days of highly controlled low energy availability in endurance-trained females. The FASEB Journal, 38(21), e70157. 10.1096/r.202401780R

Camera, D. M., Burniston, J. G., Pogson, M. A., Smiles, W. J., & Hawley, J. A. (2017). Dynamic proteome profiling of individual proteins in human skeletal muscle after a high-fat diet and resistance exercise. The FASEB Journal, 31(12), 5478–5494. 10.1096/r.201700531R

Carbone, J. W., Pasiakos, S. M., Vislocky, L. M., Anderson, J. M., & Rodriguez, N. R. (2014). Effects of short-term energy deficit on muscle protein breakdown and intramuscular proteolysis in normal-weight young adults. Applied Physiology, Nutrition, and Metabolism, 39(8), 960–968. 10.1139/apnm-2013-0433

Coen, P. M., Menshikova, E. V., Distefano, G., Zheng, D., Tanner, C. J., Standley, R. A., Helbling, N. L., Dubis, G. S., Ritov, V. B., Xie, H., Desimone, M. E., Smith, S. R., Stefanovic-Racic, M., Toledo, F. G. S., Houmard, J. A., & Goodpaster, B. H. (2015). Exercise and Weight Loss Improve Muscle Mitochondrial Respiration, Lipid Partitioning, and Insulin Sensitivity After Gastric Bypass Surgery. Diabetes, 64(11), 3737–3750. 10.2337/db15-0809

Conte, C., Hall, K. D., & Klein, S. (2024). Is Weight Loss–Induced Muscle Mass Loss Clinically Relevant? JAMA, 332(1), 9. 10.1001/jama.2024.6586

Cordeiro, A. V., Brícola, R. S., Braga, R. R., Lenhare, L., Silva, V. R. R., Anaruma, C. P., Katashima, C. K., Crisol, B. M., Simabuco, F. M., Silva, A. S. R., Cintra, D. E., Moura, L. P., Pauli, J. R., & Ropelle, E. R. (2020). Aerobic Exercise Training Induces the Mitonuclear Imbalance and UPRmt in the Skeletal Muscle of Aged Mice. The Journals of Gerontology: Series A, 75(12), 2258–2261. 10.1093/gerona/glaa059

Das, J. K., Banskota, N., Candia, J., Griswold, M. E., Orenduff, M., De Cabo, R., Corcoran, D. L., Das, S. K., De, S., Huffman, K. M., Kraus, V. B., Kraus, W. E., Martin, C. K., Racette, S. B., Redman, L. M., Schilling, B., Belsky, D. W., & Ferrucci, L. (2023). Calorie restriction modulates the transcription of genes related to stress response and longevity in human muscle: The CALERIE study. Aging Cell, 22(12), e13963. 10.1111/acel.13963

Davis, M. E., Gumucio, J. P., Sugg, K. B., Bedi, A., & Mendias, C. L. (2013). MMP inhibition as a potential method to augment the healing of skeletal muscle and tendon extracellular matrix. Journal of Applied Physiology, 115(6), 884–891. 10.1152/japplphysiol.00137.2013

Doncheva, N. T., Morris, J. H., Gorodkin, J., & Jensen, L. J. (2019). Cytoscape StringApp: Network Analysis and Visualization of Proteomics Data. Journal of Proteome Research, 18(2), 623–632. 10.1021/acs.jproteome.8b00702

Draicchio, F., Behrends, V., Tillin, N. A., Hurren, N. M., Sylow, L., & Mackenzie, R. (2022). Involvement of the extracellular matrix and integrin signalling proteins in skeletal muscle glucose uptake. The Journal of Physiology, 600(20), 4393–4408. 10.1113/JP283039

Eden, E., Navon, R., Steinfeld, I., Lipson, D., & Yakhini, Z. (2009). GOrilla: A tool for discovery and visualization of enriched GO terms in ranked gene lists. BMC Bioinformatics, 10(1), 48. 10.1186/1471-2105-10-48

Evans, W. J., & Lexell, J. (1995). Human Aging, Muscle Mass, and Fiber Type Composition. The Journals of Gerontology Series A: Biological Sciences and Medical Sciences, 50A(Special), 11–16. 10.1093/gerona/50A.Special_Issue.11

Granata, C., Caruana, N. J., Botella, J., Jamnick, N. A., Huynh, K., Kuang, J., Janssen, H. A., Reljic, B., Mellett, N. A., Laskowski, A., Stait, T. L., Frazier, A. E., Coughlan, M. T., Meikle, P. J., Thorburn, D. R., Stroud, D. A., & Bishop, D. J. (2021). High-intensity training induces non-stoichiometric changes in the mitochondrial proteome of human skeletal muscle without reorganisation of respiratory chain content. Nature Communications, 12(1), 7056. 10.1038/s41467-021-27153-3

Hesketh, S. J., Sutherland, H., Lisboa, P. J., Jarvis, J. C., & Burniston, J. G. (2020). Adaptation of rat fast-twitch muscle to endurance activity is underpinned by changes to protein degradation as well as protein synthesis. The FASEB Journal, 34(8), 10398–10417. 10.1096/r.202000668RR

Huang, J., Gong, Z., Ghosal, G., & Chen, J. (2009). SOSS complexes participate in the maintenance of genomic stability. Molecular Cell, 35(3), 384–393. 10.1016/j.molcel.2009.06.011

Iraki, J., Paulsen, G., Garthe, I., Slater, G., & Areta, J. L. (2021). Reliability of resting metabolic rate between and within day measurements using the Vyntus CPX system and comparison against predictive formulas. Nutrition and Health, 026010602110573. 10.1177/02601060211057324

Jasienska, G. (2003). Energy Metabolism and the Evolution of Reproductive Suppression in the Human Female. Acta Biotheoretica, 51(1), 1–18. 10.1023/A:1023035321162

Jasienska, G., Bribiescas, R. G., Furberg, A.-S., Helle, S., & Núñez-de La Mora, A. (2017). Human reproduction and health: An evolutionary perspective. The Lancet, 390(10093), 510–520. 10.1016/S0140-6736(17)30573-1

Jeukendrup, A. E., Areta, J. L., Van Genechten, L., Langan-Evans, C., Pedlar, C. R., Rodas, G., Sale, C., & Walsh, N. P. (2024). Does Relative Energy Deficiency in Sport (REDs) Syndrome Exist? Sports Medicine. 10.1007/s40279-024-02108-y

Jeukendrup, A. E., & Wallis, G. A. (2005). Measurement of Substrate Oxidation During Exercise by Means of Gas Exchange Measurements. International Journal of Sports Medicine, 26, S28–S37. 10.1055/s-2004-830512

Krishnan, A., Li, X., Kao, W.-Y., Viker, K., Butters, K., Masuoka, H., Knudsen, B., Gores, G., & Charlton, M. (2012). Lumican, an extracellular matrix proteoglycan, is a novel requisite for hepatic fibrosis. Laboratory Investigation; a Journal of Technical Methods and Pathology, 92(12), 1712–1725. 10.1038/labinvest.2012.121

Kumar, L., & Futschik, M. E. (2007). Mfuzz: A software package for soft clustering of microarray data. Bioinformation, 2(1), 5–7. 10.6026/97320630002005

Lee, J. Y., Park, S. J., Kim, D. A., Lee, S. H., Koh, J.-M., & Kim, B.-J. (2020). Muscle-Derived Lumican Stimulates Bone Formation via Integrin α2β1 and the Downstream ERK Signal. Frontiers in Cell and Developmental Biology, 8, 565826. 10.3389/fcell.2020.565826

Legeay, M., Doncheva, N. T., Morris, J. H., & Jensen, L. J. (2020). Visualize omics data on networks with Omics Visualizer, a Cytoscape App. F1000Research, 9, 157. 10.12688/f1000research.22280.2

Lenth, R. (2025). Emmeans: Estimated Marginal Means, aka Least-Squares Means. R package version 1.10. https://rvlenth.github.io/emmeans/

Loucks, A. B., Verdun, M., Heath, E. M., & (With the Technical Assistance of T. Law, Sr. and J. R. Thuma). (1998). Low energy availability, not stress of exercise, alters LH pulsatility in exercising women. Journal of Applied Physiology, 84(1), 37–46. 10.1152/jappl.1998.84.1.37

Magkos, F., Fraterrigo, G., Yoshino, J., Luecking, C., Kirbach, K., Kelly, S. C., de las Fuentes, L., He, S., Okunade, A. L., Patterson, B. W., & Klein, S. (2016). Effects of Moderate and Subsequent Progressive Weight Loss on Metabolic Function and Adipose Tissue Biology in Humans with Obesity. Cell Metabolism, 23(4), 591–601. 10.1016/j.cmet.2016.02.005

Mahdy, M. A. A. (2019). Skeletal muscle fibrosis: An overview. Cell and Tissue Research, 375(3), 575–588. 10.1007/s00441-018-2955-2

Marosi, K., Moehl, K., Navas-Enamorado, I., Mitchell, S. J., Zhang, Y., Lehrmann, E., Aon, M. A., Cortassa, S., Becker, K. G., & Mattson, M. P. (2018). Metabolic and molecular framework for the enhancement of endurance by intermittent food deprivation. The FASEB Journal, r.201701378RR. 10.1096/r.201701378RR

McCabe, B. J., Bederman, I. R., Croniger, C., Millward, C., Norment, C., & Previs, S. F. (2006). Reproducibility of gas chromatography-mass spectrometry measurements of 2H labeling of water: Application for measuring body composition in mice. Analytical Biochemistry, 350(2), 171–176. 10.1016/j.ab.2006.01.020

McKiernan, S. H., Colman, R. J., Aiken, E., Evans, T. D., Beasley, T. M., Aiken, J. M., Weindruch, R., & Anderson, R. M. (2012). Cellular adaptation contributes to calorie restriction-induced preservation of skeletal muscle in aged rhesus monkeys. Experimental Gerontology, 47(3), 229–236. 10.1016/j.exger.2011.12.009

Melouane, A., Yoshioka, M., & St-Amand, J. (2020). Extracellular matrix/mitochondria pathway: A novel potential target for sarcopenia. Mitochondrion, 50, 63–70. 10.1016/j.mito.2019.10.007

Morin, E., & Winterhalder, B. (2024). Ethnography and ethnohistory support the efficiency of hunting through endurance running in humans. Nature Human Behaviour. 10.1038/s41562-024-01876-x

Müller, M. J., Enderle, J., Pourhassan, M., Braun, W., Eggeling, B., Lagerpusch, M., Glüer, C.-C., Kehayias, J. J., Kiosz, D., & Bosy-Westphal, A. (2015). Metabolic adaptation to caloric restriction and subsequent refeeding: The Minnesota Starvation Experiment revisited. The American Journal of Clinical Nutrition, 102(4), 807–819. 10.3945/ajcn.115.109173

Nana, A., Slater, G. J., Hopkins, W. G., Halson, S. L., Martin, D. T., West, N. P., & Burke, L. M. (2016). Importance of Standardized DXA Protocol for Assessing Physique Changes in Athletes. International Journal of Sport Nutrition and Exercise Metabolism, 26(3), 259–267. 10.1123/ijsnem.2013-0111

Newell, M. L., Hunter, A. M., Lawrence, C., Tipton, K. D., & Galloway, S. D. R. (2015). The Ingestion of 39 or 64 g·hr−1 of Carbohydrate is Equally Effective at Improving Endurance Exercise Performance in Cyclists. International Journal of Sport Nutrition and Exercise Metabolism, 25(3), 285–292. 10.1123/ijsnem.2014-0134

Nishimura, Y., Bittel, A. J., Stead, C. A., Chen, Y.-W., & Burniston, J. G. (2023). Facioscapulohumeral Muscular Dystrophy is Associated With Altered Myoblast Proteome Dynamics. Molecular & Cellular Proteomics, 22(8), 100605. 10.1016/j.mcpro.2023.100605

Oxfeldt, M., Phillips, S. M., Andersen, O. E., Johansen, F. T., Bangshaab, M., Risikesan, J., McKendry, J., Melin, A. K., & Hansen, M. (2023). Low energy availability reduces myofibrillar and sarcoplasmic muscle protein synthesis in trained females. The Journal of Physiology, JP284967. 10.1113/JP284967

Pasiakos, S. M., Vislocky, L. M., Carbone, J. W., Altieri, N., Konopelski, K., Freake, H. C., Anderson, J. M., Ferrando, A. A., Wolfe, R. R., & Rodriguez, N. R. (2010). Acute Energy Deprivation Affects Skeletal Muscle Protein Synthesis and Associated Intracellular Signaling Proteins in Physically Active Adults. The Journal of Nutrition, 140(4), 745–751. 10.3945/jn.109.118372

Perez-Riverol, Y., Csordas, A., Bai, J., Bernal-Llinares, M., Hewapathirana, S., Kundu, D. J., Inuganti, A., Griss, J., Mayer, G., Eisenacher, M., Pérez, E., Uszkoreit, J., Pfeuffer, J., Sachsenberg, T., Yılmaz, Ş., Tiwary, S., Cox, J., Audain, E., Walzer, M., … Vizcaíno, J. A. (2019). The PRIDE database and related tools and resources in 2019: Improving support for quantification data. Nucleic Acids Research, 47(D1), D442–D450. 10.1093/nar/gky1106

Pietzner, M., Uluvar, B., Kolnes, K. J., Jeppesen, P. B., Frivold, S. V., Skattebo, Ø., Johansen, E. I., Skålhegg, B. S., Wojtaszewski, J. F. P., Kolnes, A. J., Yeo, G. S. H., O’Rahilly, S., Jensen, J., & Langenberg, C. (2024). Systemic proteome adaptions to 7-day complete caloric restriction in humans. Nature Metabolism. 10.1038/s42255-024-01008-9

Pontzer, H., & McGrosky, A. (2022). Balancing growth, reproduction, maintenance, and activity in evolved energy economies. Current Biology, 32(12), R709–R719. 10.1016/j.cub.2022.05.018

Pourteymour, S., Lee, S., Langleite, T. M., Eckardt, K., Hjorth, M., Bindesbøll, C., Dalen, K. T., Birkeland, K. I., Drevon, C. A., Holen, T., & Norheim, F. (2015). Perilipin 4 in human skeletal muscle: Localization and effect of physical activity. Physiological Reports, 3(8), e12481. 10.14814/phy2.12481

Prado, C. M., Phillips, S. M., Gonzalez, M. C., & Heymsfield, S. B. (2024). Muscle matters: The effects of medically induced weight loss on skeletal muscle. The Lancet Diabetes & Endocrinology, S2213858724002729. 10.1016/S2213-8587(24)00272-9

Prentice, A. M. (2005). Starvation in humans: Evolutionary background and contemporary implications. Mechanisms of Ageing and Development, 126(9), 976–981. 10.1016/j.mad.2005.03.018

Rhoads, T. W., Clark, J. P., Gustafson, G. E., Miller, K. N., Conklin, M. W., DeMuth, T. M., Berres, M. E., Eliceiri, K. W., Vaughan, L. K., Lary, C. W., Beasley, T. M., Colman, R. J., & Anderson, R. M. (2020). Molecular and Functional Networks Linked to Sarcopenia Prevention by Caloric Restriction in Rhesus Monkeys. Cell Systems, S2405471219304636. 10.1016/j.cels.2019.12.002

Rosenbaum, M., Goldsmith, R. L., Haddad, F., Baldwin, K. M., Smiley, R., Gallagher, D., & Leibel, R. L. (2018). Triiodothyronine and leptin repletion in humans similarly reverse weight-loss-induced changes in skeletal muscle. American Journal of Physiology-Endocrinology and Metabolism, 315(5), E771–E779. 10.1152/ajpendo.00116.2018

Sharma, V., Eckels, J., Taylor, G. K., Shulman, N. J., Stergachis, A. B., Joyner, S. A., Yan, P., Whiteaker, J. R., Halusa, G. N., Schilling, B., Gibson, B. W., Colangelo, C. M., Paulovich, A. G., Carr, S. A., Jaffe, J. D., MacCoss, M. J., & MacLean, B. (2014). Panorama: A Targeted Proteomics Knowledge Base. Journal of Proteome Research, 13(9), 4205–4210. 10.1021/pr5006636

Shepherd, S. O., Cocks, M., Meikle, P. J., Mellett, N. A., Ranasinghe, A. M., Barker, T. A., Wagenmakers, A. J. M., & Shaw, C. S. (2017). Lipid droplet remodelling and reduced muscle ceramides following sprint interval and moderate-intensity continuous exercise training in obese males. International Journal of Obesity, 41(12), 1745–1754. 10.1038/ijo.2017.170

Shepherd, S. O., Cocks, M., Tipton, K. D., Ranasinghe, A. M., Barker, T. A., Burniston, J. G., Wagenmakers, A. J. M., & Shaw, C. S. (2013). Sprint interval and traditional endurance training increase net intramuscular triglyceride breakdown and expression of perilipin 2 and 5. The Journal of Physiology, 591(3), 657–675. 10.1113/jphysiol.2012.240952

Shepherd, S. O., Strauss, J. A., Wang, Q., Dube, J. J., Goodpaster, B., Mashek, D. G., & Chow, L. S. (2017). Training alters the distribution of perilipin proteins in muscle following acute free fatty acid exposure. The Journal of Physiology, 595(16), 5587–5601. 10.1113/JP274374

Speakman, J. (2007). A Nonadaptive Scenario Explaining the Genetic Predisposition to Obesity: The “Predation Release” Hypothesis. Cell Metabolism, 6(1), 5–12. 10.1016/j.cmet.2007.06.004

Speakman, J. R., & Mitchell, S. E. (2011). Caloric restriction. Molecular Aspects of Medicine, 32(3), 159–221. 10.1016/j.mam.2011.07.001

Storey, J. D., & Tibshirani, R. (2003). Statistical significance for genomewide studies. Proceedings of the National Academy of Sciences, 100(16), 9440–9445. 10.1073/pnas.1530509100

Toledo, F. G. S., Menshikova, E. V., Azuma, K., Radiková, Z., Kelley, C. A., Ritov, V. B., & Kelley, D. E. (2008). Mitochondrial Capacity in Skeletal Muscle Is Not Stimulated by Weight Loss Despite Increases in Insulin Action and Decreases in Intramyocellular Lipid Content. Diabetes, 57(4), 987–994. 10.2337/db07-1429

Toledo, F. G. S., Menshikova, E. V., Ritov, V. B., Azuma, K., Radikova, Z., DeLany, J., & Kelley, D. E. (2007). Effects of Physical Activity and Weight Loss on Skeletal Muscle Mitochondria and Relationship With Glucose Control in Type 2 Diabetes. Diabetes, 56(8), 2142–2147. 10.2337/db07-0141

Toledo, F. G. S., Watkins, S., & Kelley, D. E. (2006). Changes Induced by Physical Activity and Weight Loss in the Morphology of Intermyofibrillar Mitochondria in Obese Men and Women. The Journal of Clinical Endocrinology & Metabolism, 91(8), 3224–3227. 10.1210/jc.2006-0002

Ubaida-Mohien, C., Lyashkov, A., Gonzalez-Freire, M., Tharakan, R., Shardell, M., Moaddel, R., Semba, R. D., Chia, C. W., Gorospe, M., Sen, R., & Ferrucci, L. (2019). Discovery proteomics in aging human skeletal muscle finds change in spliceosome, immunity, proteostasis and mitochondria. eLife, 8, e49874. 10.7554/eLife.49874

Van Rosmalen, L., Zhu, J., Maier, G., Gacasan, E. G., Lin, T., Zhemchuzhnikova, E., Rothenberg, V., Razu, S., Deota, S., Ramasamy, R. K., Sah, R. L., McCulloch, A. D., Hut, R. A., & Panda, S. (2024). Multi-organ transcriptome atlas of a mouse model of relative energy deficiency in sport. Cell Metabolism, 36(9), 2015–2037.e6. 10.1016/j.cmet.2024.08.001

Wang, H., Sreenivasan, U., Hu, H., Saladino, A., Polster, B. M., Lund, L. M., Gong, D., Stanley, W. C., & Sztalryd, C. (2011). Perilipin 5, a lipid droplet-associated protein, provides physical and metabolic linkage to mitochondria. Journal of Lipid Research, 52(12), 2159–2168. 10.1194/jlr.M017939

Wickham, H., & Sievert, C. (2009). ggplot2: Elegant graphics for data analysis (Vol. 10). springer New York.

Williams, A. S., Crown, S. B., Lyons, S. P., Koves, T. R., Wilson, R. J., Johnson, J. M., Slentz, D. H., Kelly, D. P., Grimsrud, P. A., Zhang, G.-F., & Muoio, D. M. (2024). Ketone flux through BDH1 supports metabolic remodeling of skeletal and cardiac muscles in response to intermittent time-restricted feeding. Cell Metabolism, 36(2), 422–437.e8. 10.1016/j.cmet.2024.01.007

Wolins, N. E., Brasaemle, D. L., & Bickel, P. E. (2006). A proposed model of fat packaging by exchangeable lipid droplet proteins. FEBS Letters, 580(23), 5484–5491. 10.1016/j.febslet.2006.08.040

Yang, L., Licastro, D., Cava, E., Veronese, N., Spelta, F., Rizza, W., Bertozzi, B., Villareal, D. T., Hotamisligil, G. S., Holloszy, J. O., & Fontana, L. (2016). Long-Term Calorie Restriction Enhances Cellular Quality-Control Processes in Human Skeletal Muscle. Cell Reports, 14(3), 422–428. 10.1016/j.celrep.2015.12.042

Zanini, G., Selleri, V., Malerba, M., Solodka, K., Sinigaglia, G., Nasi, M., Mattioli, A. V., & Pinti, M. (2023). The Role of Lonp1 on Mitochondrial Functions during Cardiovascular and Muscular Diseases. Antioxidants, 12(3), 598. 10.3390/antiox12030598

Zhang, B., Kirov, S., & Snoddy, J. (2005). WebGestalt: An integrated system for exploring gene sets in various biological contexts. Nucleic Acids Research, 33(Web Server), W741–W748. 10.1093/nar/gki475

Zhang, Q., Zheng, J., Qiu, J., Wu, X., Xu, Y., Shen, W., & Sun, M. (2017). ALDH2 restores exhaustive exercise-induced mitochondrial dysfunction in skeletal muscle. Biochemical and Biophysical Research Communications, 485(4), 753–760. 10.1016/j.bbrc.2017.02.124

