## Supplementary material for "Skeletal muscle proteomic responses to energy deficit with concomitant aerobic exercise in humans": Protocol

***Effects of Acute Energy Deficit with Concomitant Aerobic Exercise on Skeletal Muscle Quality***

Weight-loss strategies that use energy restriction alone can lead to impaired muscle mass, which can also further impair health of individuals with metabolic conditions, such as type 2 diabetics. Skeletal muscle mass and function are key to maintaining a healthy metabolism and quality of life throughout the lifespan. The combination of exercise and calorie restriction is a powerful intervention for reducing body weight and improving the metabolism and health status of healthy, overweight and obese individuals. We have previously shown that skeletal muscle protein synthesis is reduced with energy restriction and that resistance-type exercise can reverse this negative effect. Aerobic-type exercise also has the capacity to stimulate muscle protein synthesis and improve the quality of muscle by increasing mitochondrial protein synthesis. Nonetheless, how energy deficit overlayed on top of aerobic exercise modulates skeletal muscle quality is not well characterised. Furthermore, existing studies have looked only into mixed (unspecific) protein synthesis of different intracellular compartments without providing details on how protein synthesis for specific proteins is regulated.

This project will investigate the mechanisms behind the effect of energy restriction and concomitant aerobic exercise on skeletal muscle quality through the use of dynamic proteomic profiling.

Aerobic exercise alone is a well-established intervention to increase mitochondrial capacity and skeletal muscle function, but the effect on skeletal muscle of overlaying energy restriction while performing aerobic exercise is not well characterised. Recent findings in Rhesus monkeys, has shown that life-long caloric restriction has a positive effect on skeletal muscle. These findings show that caloric restriction not only maintains contractile content of muscle, but also rescues the age-related decline of skeletal muscle mitochondrial content and capacity. However, physical activity in that study was not controlled and appeared to be higher in the caloric restriction group, representing an important confounding factor (Rhoads et al., 2020). The proposed study herein will include a short period of tightly controlled exercise and dietary intake to investigate the effect of energy deficit while performing aerobic exercise.

Objectives & Hypothesis:

The primary research question associated with this study is: 'What is the combined effect of an energy deficit and aerobic exercise training on muscle quality (abundance and synthesis rates of individual, sarcoplasmic and mitochondrial proteins) over a five-day period in healthy males?'

We hypothesize that energy deficit will increase skeletal muscle mitochondrial proteins while individuals’ endocrine and metabolic parameters are consistent with a response for energy preservation.

***OUTCOMES***

*Primary outcomes:*

*1. Skeletal muscle proteomics (dynamic proteomic profiling).*

*Secondary outcomes:*

1. *Body mass.*
2. *Body composition (DXA & BIA).*
3. *Blood concentration of metabolites.*
4. *Blood concentration of hormones.*
5. *Skeletal muscle glycogen.*
6. *Skeletal muscle lipid droplets.*
7. *Resting Metabolic Rate.*
8. *Substrate use during exercise.*

***DESIGN, PARTICIPANTS AND SAMPLE AND DATA COLLECTION METHODOLOGY***

A schematic overview of the experimental protocol is shown in Figure 1.

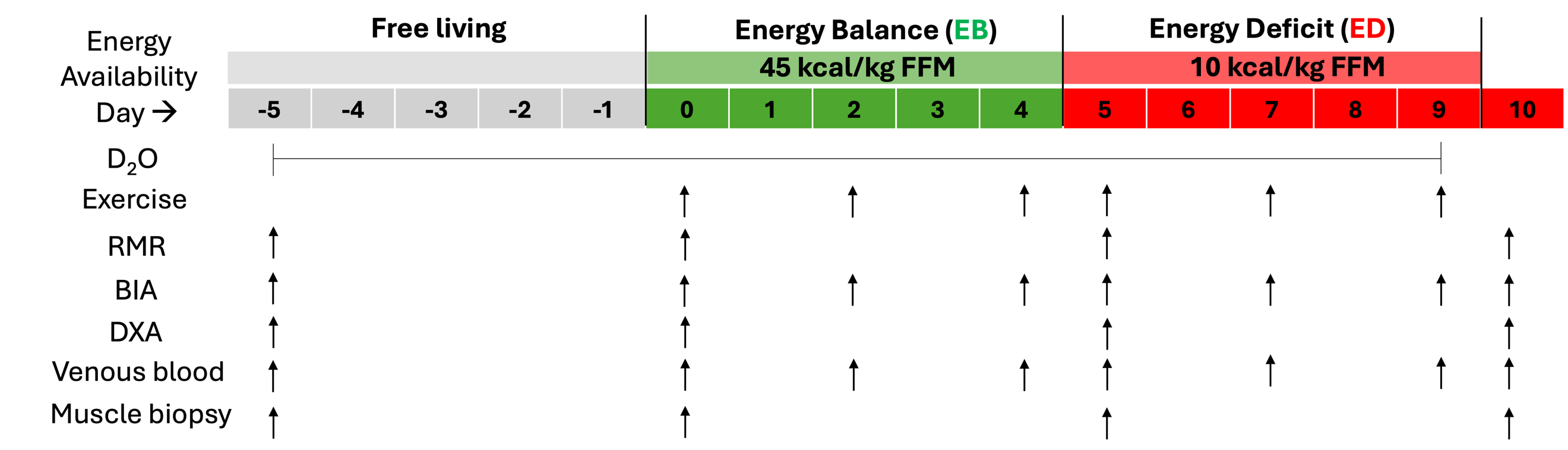

**Figure 1.** Schematic overview of experimental protocol including provision of D_2_O, laboratory-based exercise, assessment of resting metabolic rate (RMR), body composition (through dual x-ray absorptiometry (DXA) and bioimpedance analysis (BIA), collection of blood, saliva and skeletal muscle.

***Study inclusion criteria:***

- *Gender/Sex, Male*
- *Age, 18 – 40 years*
- *Body fat, ~18 – 26%*
- *Health, healthy, as determined by pre-participation questionnaires*
- *Training Status, Regularly Exercising/Aerobically trained (3-4 aerobic training sessions/week, 3-5 hrs/week)*
- *Non-smoker*
- *Weight-stable (within 2 kg) for the past 6-months (self-reported)*
- *Eating attitude test (EAT-26) score < 20*

**Exclusion Criteria:**

- Gender/Sex, Female/Other
- Age, <18 - >40 years
- Health, deemed unable to perform exercise (assessed via readiness to exercise questionnaire)
- Medical Condition, those with any previous diagnosis of: Osteoporosis/low bone mineral density, cardio-vascular disease, diabetes mellitus, cerebrovascular disease, blood-related illness/disorder, asthma or other respiratory illness/disorder, liver disease, kidney disease, gastrointestinal disease, eating disorder or disordered eating
- Those currently taking prescription medication or unwell with a cold or virus at the time of participation.
- Those unwilling to adhere to the study's methodological requirements (including adhering to alterations in diet and training – including alcohol abstention) from the day prior to intervention onset (24 hrs pre-intervention) to completion of follow-up assessments (day 10).
- Those following a restrictive diet (e.g. vegetarians/vegans)
- Those with food allergy/intolerances incompatible with study diets
- Training status, Does not train aerobically 3 + times/week (over past 6 months on average)
- Current smoker

**Baseline anthropometric and fitness testing:**

Participants attended the laboratory for a baseline test at least two days, and no more than seven days prior to their proposed start-date for the study intervention phase. The participant voided their bladder and body composition was assessed with Bio-impedance analysis (BIA)(Bosy-Westphal et al., 2017).

Participants were fitted with a Bluetooth heart rate (HR) monitor (Polar H7, Polar, Kempele, Finland) and set up on an electro-magnetically braked cycle ergometer (Lode Corival cpet: Lode, Groningen, Netherlands). Participants then performed a two-part incremental cycle test to determine sub-maximal (V̇O2) and peak power output/maximal oxygen consumption (V̇O_2max_ Moxus Modular Metabolic System, AEI Technologies, Pittsburgh, PA, USA), lactate threshold (LT; Biosen C-Line, EKF Diagnostic, Cardiff, UK) (Newell et al., 2015) and followed by a brief 20- minute cycling familiarisation session replicating the demands of laboratory-based exercise.

**Resting Metabolic Rate (RMR):**

To assess RMR, participants lay supine and quietly on a medical bed in a dimly lit room for five minutes. A transparent ventilated hood was then placed over the participant’s head connecting with an open circuit indirect calorimeter (GEMNutrition Ltd., Warrington, UK). The participant was instructed to relax, lay as still as possible, and to breathe normally over the duration of the assessment. The RMR assessment was conducted over a further 25 minutes, with only the final 20 minutes of data used for data analysis (Iraki et al., 2021). The 20-minute average V̇O_2_ and V̇CO_2_, was used to calculate the average resting energy expenditure (kcal•min^-1^) over the assessment period and extrapolated to reflect a 24 hour (daily) resting metabolic rate (kcal•day^-1^).

**Muscle biopsy:**

Skeletal Muscle biopsies were taken from the vastus lateralis portion of the quadriceps muscle using the Using the Weil-Blakesley Conchotome technique in the morning after RMR assessment. A single muscle biopsy was collected at each time-point, each day, from a separate incision site which was 2-3 cm from any previous biopsy sites.

**Venous blood sampling:**

Participants had a fasted morning venous blood sample drawn from the ante-cubital vein by a trained phlebotomist. Approximately 18 – 26 ml of blood was drawn at each time-point. Samples were drawn into singular 6 ml whole blood (serum), 8 ml serum separator tube (SST), 6 ml ethylenediaminetetraacetic acid (EDTA), and 6 ml lithium heparin (LHep) vacutainers (BD Vacutainer, Becton, Dickinson and Company, Franklin Lakes, NJ, USA). All vacutainers were inverted eight-times following collection, in line with our laboratory guidelines. Plasma samples (EDTA and LHep) were placed on ice immediately, whilst serum and SST vacutainers were allowed to clot at room temperature for 30 minutes, before being placed on ice until centrifugation. Blood samples were centrifuged for 10-minutes at 4˚C and a relative centrifugal force of 1200g, plasma and serum were then stored at -80˚C for subsequent analysis.

**D_2_O provision:**

After RMR measurement, and sampling of blood and skeletal muscle biopsy samples on day -5, ahead of the free-living period, participants were provided with a *500 ml D_2_O pre-load and then a daily 50 ml D_2_O top-up dose until day 9 (inclusive).*

**Body composition:**

Changes in body composition during the study were assessed using dual-energy x-ray absorptiometry (DXA; QDR Series Discovery A; Hologic Inc., Marlborough Massachusetts, USA, software version 12:4:3) following standardised procedures (Nana et al., 2016).

BIA was used to assess body composition and total body water content changes using an 8-electrode BIA machine (SECA mBCA 515; SECA GMBH, Hamburg, Germany), following manufacturer guidelines and the computer-linked analysis software (SECA Analytics 115: SECA GMBH, Hamburg, Germany).

**Diet:**

Diets were designed to maintain body mass (Energy Balance; EB) or produce body mass loss (Energy Deficit; ED) through achieving an average energy availability (Loucks et al., 1998) of 45 and 10 kcal *• Fat Free Mass (FFM)•day ^-1^*, with predicted average body mass losses of 0 and ~3 kg (Areta et al., 2021), respectively.

Diets were pre-packaged custom-made to provide a daily energy intake of 54 and 19 kcal *• Fat Free Mass (FFM) • day ^-1^* in EB and ED, respectively with 60, 20 and 20% of energy derived from carbohydrates, fat and protein, respectively, regardless of the dietary intervention, and ~60% of energy derived from white potatoes. Details of diets consumed are shown in supplementary table 2.

**Laboratory-based exercise and exercise indirect calorimetry:**

Cycling ergometry exercise sessions were completed following the collection of all measurements (RMR, DXA and BIA) and samples (blood and muscle).

Exercise was initiated after fitting a mouthpiece and nose-clip for indirect calorimetry with four consecutive three-minute stages of progressive workload at 50 W, 75 W, 100 W and 125 W to provide a standardised warm-up and for assessment of sub-maximal cycling efficiency and substrate utilisation. After the standardised 12-minute warm-up, workload was set to the power output corresponding to 60% V̇O_2max_.

Participants kept the mouthpiece and nose clip on for a further 3-minutes, up to 15-minutes of exercise. Heart rate was recorded throughout and rate of perceived exertion (RPE) (Borg, 1982) was recorded in the final minute of each stage and substrate oxidation and net exercise energy expenditure was calculated from the mean values of expired O_2_ and CO_2_ from the final 60 seconds of each stage.

Thereafter, participants were asked to insert the mouthpiece and fit the nose clip for the final 2.5 minutes of every 15 minutes of exercise, to allow the collection of two minutes of expired gas samples for every 15 mins of exercise (e.g., from 27.5 – 30 mins) assessment of substrate utilisation. HR and RPE were recorded in the final minute of each 15-minute exercise block, with substrate oxidation and net exercise energy expenditure calculated from the mean values of expired O_2_ and CO_2_ from the final 60s of each stage. Participants cycled until achieving net exercise energy expenditure equating to 15 kcal • kg FFM^-1^, calculated in real time from indirect calorimetry. Every 30 minutes of exercise, participants stopped exercising for a 5-minute rest period.

To avoid hypoglycaemia during fasted exercise, participants were provided with a small meal (139 kcal; 30 g CHO, <1 g FAT, 3 g PRO) 15 min before exercise and in the 5 min rest period after the 30 and 60 min blocks of exercise.

Details of exercise related parameters are shown in supplementary table 4.

**References:**

Areta, J. L., Taylor, H. L., & Koehler, K. (2021). Low energy availability: History, definition and evidence of its endocrine, metabolic and physiological effects in prospective studies in females and males. *European Journal of Applied Physiology*, *121*(1), 1–21. https://doi.org/10.1007/s00421-020-04516-0

Borg, G. A. (1982). Psychophysical bases of perceived exertion. *Med Sci Sports Exerc*, *14*(5), 377–381.

Bosy-Westphal, A., Jensen, B., Braun, W., Pourhassan, M., Gallagher, D., & Müller, M. J. (2017). Quantification of whole-body and segmental skeletal muscle mass using phase-sensitive 8-electrode medical bioelectrical impedance devices. *European Journal of Clinical Nutrition*, *71*(9), 1061–1067. https://doi.org/10.1038/ejcn.2017.27

Iraki, J., Paulsen, G., Garthe, I., Slater, G., & Areta, J. L. (2021). Reliability of resting metabolic rate between and within day measurements using the Vyntus CPX system and comparison against predictive formulas. *Nutrition and Health*, 026010602110573. https://doi.org/10.1177/02601060211057324

Loucks, A. B., Verdun, M., Heath, E. M., & (With the Technical Assistance of T. Law, Sr. and J. R. Thuma). (1998). Low energy availability, not stress of exercise, alters LH pulsatility in exercising women. *Journal of Applied Physiology*, *84*(1), 37–46. https://doi.org/10.1152/jappl.1998.84.1.37

Nana, A., Slater, G. J., Hopkins, W. G., Halson, S. L., Martin, D. T., West, N. P., & Burke, L. M. (2016). Importance of Standardized DXA Protocol for Assessing Physique Changes in Athletes. *International Journal of Sport Nutrition and Exercise Metabolism*, *26*(3), 259–267. https://doi.org/10.1123/ijsnem.2013-0111

Newell, M. L., Hunter, A. M., Lawrence, C., Tipton, K. D., & Galloway, S. D. R. (2015). The Ingestion of 39 or 64 g·hr−1 of Carbohydrate is Equally Effective at Improving Endurance Exercise Performance in Cyclists. *International Journal of Sport Nutrition and Exercise Metabolism*, *25*(3), 285–292. https://doi.org/10.1123/ijsnem.2014-0134

Rhoads, T. W., Clark, J. P., Gustafson, G. E., Miller, K. N., Conklin, M. W., DeMuth, T. M., Berres, M. E., Eliceiri, K. W., Vaughan, L. K., Lary, C. W., Beasley, T. M., Colman, R. J., & Anderson, R. M. (2020). Molecular and Functional Networks Linked to Sarcopenia Prevention by Caloric Restriction in Rhesus Monkeys. *Cell Systems*, S2405471219304636. https://doi.org/10.1016/j.cels.2019.12.002
